# A Phylogenetic Profiling Approach Identifies Novel Ciliogenesis Genes In *Drosophila* And *C. elegans*

**DOI:** 10.1101/2022.12.28.522111

**Authors:** Jeroen Dobbelaere, Tiffany Y. Su, Balazs Erdi, Alexander Schleiffer, Alexander Dammermann

**Author notes:** Contributed equally (Jeroen Dobbelaere, Tiffany Y. Su, Balazs Erdi). Correspondence Corresponding Author: Alexander Dammermann Max Perutz Labs, Dr Bohr-Gasse 9, A-1030 Vienna, Austria.

## Abstract

Cilia are cellular projections that perform sensory and motile functions in eukaryotic cells. A defining feature of cilia is that they are evolutionarily ancient yet not universally conserved. In this study we have used the resulting presence and absence pattern in the genomes of diverse eukaryotes to identify a set of 386 human genes associated with cilium assembly or motility. Comprehensive tissue-specific RNAi in *Drosophila* and mutant analysis in *C. elegans* revealed signature ciliary defects for 70-80% of novel genes, a percentage similar to that for known genes within the cluster. Further characterization identified different phenotypic classes, including a set of genes related to the cartwheel component Bld10/Cep135 and two highly conserved regulators of cilium biogenesis. We believe this dataset to define the core set of genes required for cilium assembly and motility across eukaryotes, an invaluable resource for future studies of cilium biology and associated disorders.

## INTRODUCTION

The highly stereotypical architecture of cilia and flagella (Fig. 1A), terms often used interchangeably, can be found in all branches of the eukaryotic tree of life, albeit not in every species. Indeed, the number of cilia per cell was the defining characteristic to divide eukaryotes into two superclades, unikonts (literally: single flagellum) and bikonts (two) (Cavalier-Smith, 2002). Unikonts comprise the amoebozoa and opisthokonta (posterior flagellum), a clade that contains metazoa, fungi and choanoflagellates, while bikonts comprise the remaining clades. Unikonts typically have only one cilium and associated basal body positioned at the posterior end of the cell, whereas bikonts have two, a younger, anterior, cilium and an older, posterior, cilium. Which of the two configurations represents the ancestral state is unclear (Roger and Simpson, 2009), but what is clear is that the last common ancestor of all eukaryotes harbored both centrioles and cilia, while centriole-organized centrosomes appear to be a much later opisthokont or metazoan innovation (Azimzadeh, 2014).

**Figure 1:**
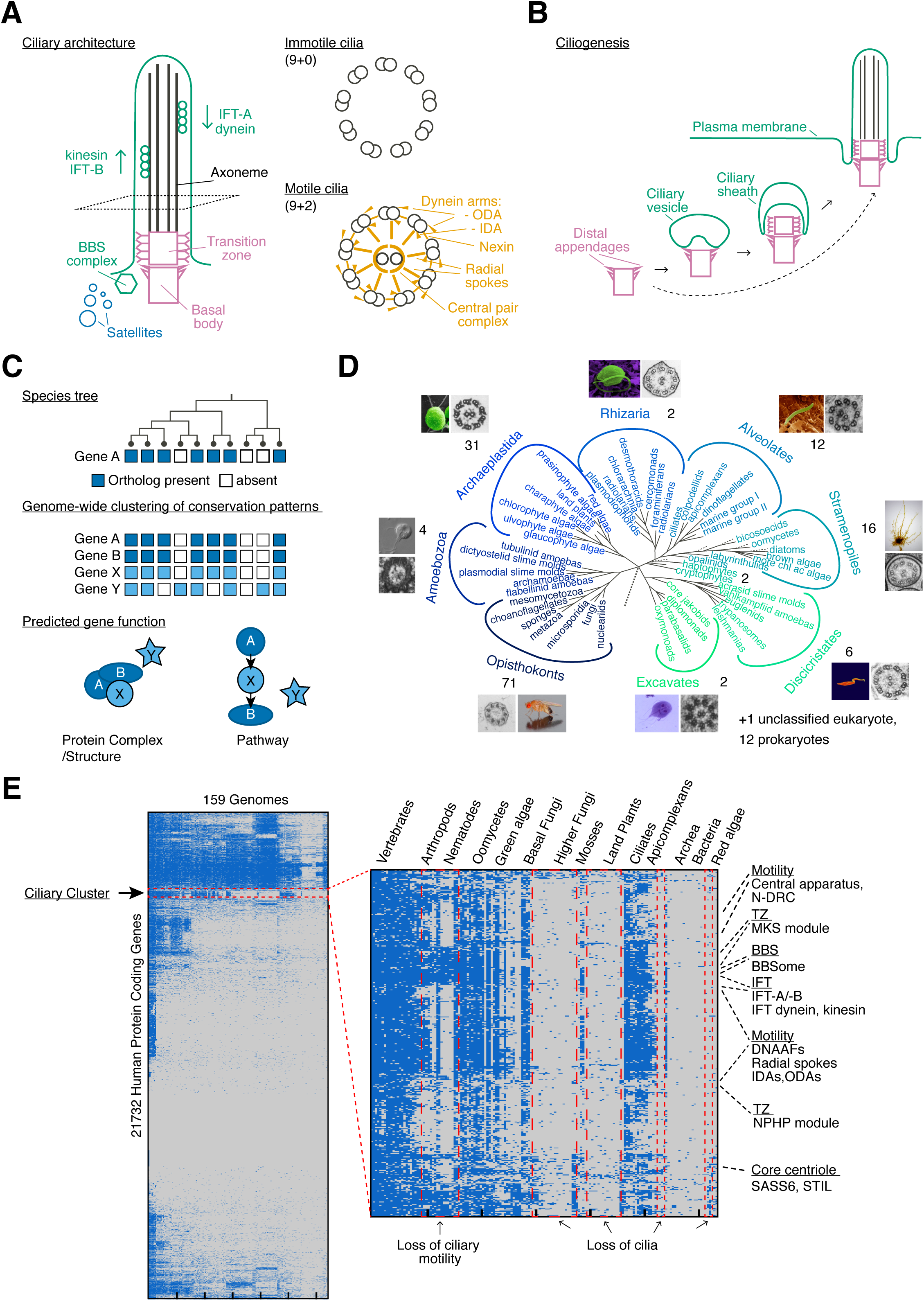
Identification of a ciliary cluster by phylogenetic profiling. (**A**) Schematic of ciliary architecture, depicting centriole-derived basal body and transition zone (pink), intraflagellar transport (IFT, green), axoneme (black) and centriolar satellites (blue). Cross sections illustrate axonemes of motile and immotile cilia. The latter display multiple additional features, including inner and outer dynein arms, nexins, radial spokes and the central pair complex (yellow). (**B**) Schematic of ciliogenesis process (based on (Reiter *et al*., 2012)). In the intracellular pathway, a vesicle forms in association with the basal body. The ciliary transition zone forms immediately distal to the basal body before the vesicle fuses with the plasma membrane, creating a ciliary pocket. Intraflagellar transport subsequently extends the ciliary axoneme beyond the transition zone. Alternatively, basal bodies dock directly to the plasma membrane prior to transition zone formation and axoneme elongation (dashed arrow). In *Plasmodium* and *Drosophila* sperm, axoneme elongation occurs in an IFT-independent manner within the cytoplasm, whereupon the cilium is extruded from the cell (not depicted). (**C**) Schematic of phylogenetic profiling approach (adapted from (Dey *et al*., 2015)). Reciprocal BLAST analysis using the human proteome as the starting point creates a presence and absence matrix for each gene across species. Genes functioning together in the same complex/structure or pathway share a similar inheritance pattern across evolution that is distinct from that of unrelated proteins, aiding in the identification of novel components (here: X). (**D**) Eukaryotic tree (Baldauf, 2008), illustrating the presence of cilia in all major groups. 147 genomes from across eukaryotes as well as 12 prokaryotic species as outgroups were chosen for our phylogenetic profiling analysis. See Table S1A. Image sources: Amoebozoa (*Phalansterium arcticum*, (Shmakova et al., 2018)), Archaeplastida (*Chlamydomonas reinhardtii*, Wolfram Weckwerth, (Huang et al., 1979)), Rhizaria (*Bigelowiella natans*, Geoff McFadden, (Moestrup and Sengco, 2001)), Alveolates (*Plasmodium falciparum*, Volker Brinkmann, (Francia et al., 2015)), Stramenopiles (*Ectocarpus siliculosus*, Delphine Scornet, (Fu et al., 2013)), Discicristates (*Trypanasoma brucei*, Philippe Bastin, (Gluenz et al., 2015)), Excavates (*Giardia lamblia*, National Institute of Infectious Diseases, Japan, (Tůmová et al., 2021)), Opisthokonts (*Drosophila melanogaster*, André Karwath, this study). (**E**) Phylogenetic profiling output, highlighting the ciliary cluster. Species nodes were manually sorted to shift ciliated species as far to the left as possible while respecting the hierarchical clustering output. Gene order is unchanged and highlights the grouping of functional subcomplexes within the cluster. See also Table S1B.

Cilia perform both sensory and motile functions. Ciliary motility is used to propel cells through fluid, as in the case of the flagellum of *Chlamydomonas* or vertebrate sperm, or fluid over the surface of the cell, as in the case of multiciliated epithelia of the vertebrate lung or oviduct. Ciliary motility usually involves dynein motor-mediated ciliary bending (Viswanadha et al., 2017), although gliding motility involving ciliary adhesion to the substrate and internal intraflagellar transport (IFT)-mediated transport of attachment sites has also been observed (Shih et al., 2013). Projecting from the cell surface and with a greatly enhanced surface-to-volume ratio, cilia are well suited to receive and transduce signals from the extracellular environment, including light, low molecular weight chemicals, proteins, and mechanical stimuli (Nachury, 2014). In vertebrates, exclusively non-motile primary cilia are found on the surface of most cell types and are involved not only in the classical senses of vision and olfaction but are also integral to many of the signaling pathways that underlie development and tissue homeostasis (Satir and Christensen, 2007). The manifold functions of cilia in vertebrates are reflected in the pleiotropic nature of human cilium-related disorders or ciliopathies, with clinical manifestations including *situs inversus*, respiratory dysfunction, infertility and hydrocephalus for disorders affecting motile cilia, while disorders affecting non-motile cilia are characterized by defects in neural tube development and limb patterning, cystic kidney, liver and pancreatic diseases, retinal degeneration, anosmia, cognitive defects and obesity (Mitchison and Valente, 2017). Yet, while the most severe ciliopathies are perinatal lethal, they nevertheless represent only the weak end of the phenotypic spectrum, with complete loss of cilia resulting in embryonic lethality in mice (Murcia et al., 2000).

During ciliogenesis, centrioles dock either to a vesicle in the cytoplasm that fuses with the plasma membrane creating a ciliary pocket (intracellular pathway) or to the plasma membrane itself (extracellular pathway) (Sorokin, 1962) (Fig. 1B). Central to this process are the transition fibers (Schmidt et al., 2012), which in vertebrates are derived from appendages present at the distal end of mature centrioles. These recruit a kinase, TTBK2, which promotes the removal of inhibitory factors present at the distal end of centrioles and the assembly of ciliary structures (Goetz et al., 2012). Formation of the outer microtubule doublets of the axoneme occurs by direct extension of centriolar microtubules, while the central pair of microtubules present in motile cilia forms *de novo* distal to the basal body (O’Toole et al., 2003). Motile cilia furthermore have multiple accessory structures decorating the axoneme (nexins, outer and inner dynein arms), which mediate ciliary bending (Ishikawa, 2017). Separating the basal body from the cilium proper is the transition zone, an elaborate structure characterized by Y-links connecting axonemal microtubules to the ciliary membrane. The transition zone serves to restrict protein access, establishing the cilium as a distinct cellular compartment (Reiter et al., 2012). Extension of the axoneme within this compartment requires microtubule motor-driven intraflagellar transport (IFT) delivering material to the ciliary tip (Kozminski et al., 1993).

Not all aspects of ciliogenesis are presently equally well understood. This applies in particular to the early steps involving docking of centrioles to the plasma membrane and initiation of transition zone assembly/axoneme extension. We have an increasingly detailed understanding of the molecular mechanisms underlying the intracellular pathway (Shakya and Westlake, 2021) in vertebrates. Yet, the degree to which these are relevant to the less well studied extracellular pathway is unclear, with key players apparently dispensable in this context and not conserved beyond metazoans (Ganga et al., 2021; Stuck et al., 2021). The role of centriolar appendages is also far from universal, with appendages seemingly lacking in *Drosophila* and *C. elegans* (Gottardo et al., 2015), without impairing basal body docking and initiation of ciliogenesis in these species. It is possible that different species have evolved different mechanisms for the same biological challenge. Alternatively, we do not yet possess the full set of conserved genes involved in the early steps of ciliogenesis. Missing genes likely also account for some of the still significant number of ciliopathy cases for which the causative genes have yet to be identified (Mitchison and Valente, 2017).

Our present compendium of ciliogenesis components (Vasquez et al., 2021) derives from a variety of different sources, including proteomics of isolated cilia in *Tetrahymena* (Smith et al., 2005) and *Chlamydomonas* (Pazour et al., 2005; Zhao et al., 2019) and genetic screens in *C. elegans* (Perkins et al., 1986), which led to the identification of much of the machinery for ciliary motility and IFT. Basal body and transition zone components in part have also been identified by proteomics (Andersen et al., 2003), but more commonly as the causative genes in human ciliopathies (Reiter and Leroux, 2017). Comprehensive genome-wide RNAi or mutant screens have as yet not been carried out except in cultured cells (Kim et al., 2010; Wheway et al., 2015). Instead, targeted approaches have been employed to enrich for ciliary genes. Comparative genomics in particular has proven very fruitful. Two landmark studies published in 2004 took advantage of the first fully sequenced eukaryotic genomes to define ciliary genes by simple subtractive analysis (i.e. ciliated vs. non-ciliated species), identifying key components of the ciliary trafficking machinery (Avidor-Reiss et al., 2004; Li et al., 2004). More recently, the availability of increasing numbers of sequenced genomes have enabled more sophisticated phylogenetic profiling analyses, using co-occurrence patterns across evolution to predict genes with a common function (Fig. 1C) (Dey and Meyer, 2015; Pellegrini et al., 1999). Here, we conducted a wider phylogenetic profiling approach, utilizing 147 genomes representing all major phyla of the eukaryotic tree to define a cluster of 386 human genes associated with the presence of cilia, including most of the known players involved in key steps of centriole assembly, cilium biogenesis and motility, as well as another 152 genes that have so far not been functionally characterized. Systematic RNAi-based analysis of novel genes in *Drosophila* as well as more targeted mutant analysis in *C. elegans* suggests that most if not all of these genes are likely cilium-related, with our downstream characterization focusing on a set of genes involved in early steps of the ciliogenesis pathway.

## RESULTS

### Identification of a ciliary cluster using a refined phylogenetic profiling approach

The present study emerged from a long-standing interest in the lab in the molecular mechanisms underlying centriole assembly and function in ciliogenesis, using the nematode *Caenorhabditis elegans* and the fruit fly *Drosophila melanogaster* as experimental models (Dammermann et al., 2004; Dammermann et al., 2009; Dobbelaere et al., 2020; Serwas et al., 2017). Given the highly conserved nature of centrioles and cilia, we hypothesized that the early stages of ciliogenesis including centriole to basal body conversion and initiation of axoneme extension must also be conserved across eukaryotes. To identify candidate genes potentially involved in these processes we decided to apply a refined phylogenetic profiling approach. The original landmark studies had demonstrated the utility of comparative genomics approaches to identify ciliary genes (Avidor-Reiss *et al*., 2004; Li *et al*., 2004). However, we noted that certain proteins such as the core centriolar components CENPJ/SAS-4, STIL/SAS-5/Ana2 and SASS6 tend to be missed in orthology inferences due to their high degree of sequence divergence making them drop below significance thresholds in rapidly evolving species such as *C. elegans* (Carvalho-Santos et al., 2010; Hodges et al., 2010). In setting out to generate our phylogenetic profiles we therefore chose a deliberately low BLASTp sequence similarity cutoff of 0.1, with bidirectional best match as a simple but robust (Kristensen et al., 2011) method to infer orthology. We further decided to use the human proteome as the starting point for our analysis, not only because it is the best annotated, but also because vertebrates display a lower rate of sequence divergence compared to other eukaryotes (Douzery et al., 2004; Peterson et al., 2004; Raible et al., 2005). Phylogenetic profiling approaches have grown increasingly complex, with recent studies aimed at identifying ciliary genes incorporating information on the ciliary status of species, their evolutionary relationship and/or training sets of verified ciliary genes, as well as iterative agglomerative clustering to identify orthogroups (Dey et al., 2015; Li et al., 2014; Nevers et al., 2017). We here opted instead for a simple hierarchical clustering approach, incorporating no information beyond the presence/absence pattern computed for each of the 21.732 human protein coding genes across the genomes of 147 diverse eukaryotes representing the different branches of the eukaryotic tree, as well as 12 prokaryotic species (9 archaea, 3 bacteria) as outgroups (see Fig. 1D and Table S1A). In selecting genomes, care was taken to avoid over-representing certain branches of the tree and to exclude incompletely sequenced/annotated genomes that would create false negatives in the phylogenetic analysis. Visual inspection of the clustering output revealed a set of 386 genes whose inheritance pattern was clearly distinct from that of other highly conserved genes (see Fig. 1E and Table S1B). Further analysis showed this set of genes to comprise key centriolar and ciliary genes, and their presence and absence pattern across eukaryotes to reflect the presence and absence of cilia in those species (see below). Such a single ciliary cluster had not been observed in previous analyses (Dey *et al*., 2015; Li *et al*., 2014; Nevers *et al*., 2017). We therefore sought to confirm the robustness of our analysis by reducing the number of species representing each branch of the tree by 25%, 50% and 75%. A clear ciliary cluster could still be identified in all cases, although the percentage of cluster genes remaining dropped below 80% with the most aggressive perturbation (277/386 genes when using 42 genomes, see Table S1). This is in line with published reports of diminishing returns in phylogenetic profiling approaches >100 genomes (Škunca and Dessimoz, 2015). We conclude that our phylogenetic profiling approach robustly identifies a set of genes putatively linked to cilia.

### *In silico* characterization of the ciliary phylogenetic cluster

We next set out to examine the 386 genes in the cluster by performing a comprehensive literature and database analysis, including any information on their putative orthologs in non-vertebrate experimental models (Table S2A). We classified each gene based on its level of functional characterization overall and any previously reported link to cilia on a scale of 0-3 (0 no information, 1 some indication of centriole/cilium-related function, 2 confirmed function, 3 clearly established molecular mechanism). We further set out to provide a short description of its proposed function, as well as any potential or confirmed ciliopathy association. At time of writing, 234 of the 386 genes in the cluster had a reported function in cilium assembly or motility (i.e. with a score of 2 or above), with the other 152 genes as yet functionally uncharacterized or not linked to cilia (Fig. 2A). How comprehensive is this list for known ciliary complexes and structures? As shown in Fig. 2B, more than 50% and in many cases nearer 100% of components reported to be associated with basal bodies (including centriolar and transition zone components), cilium assembly (IFT and BBS components) and motility (inner and outer dynein arm components and dynein assembly factors, nexins, N-DRC, radial spoke and central apparatus components and the only recently identified microtubule inner proteins (MIPs)) can be found in the cluster, indicating essentially complete coverage of core components. Notably missing are components linked to centrosomes (PCM and centriolar satellites), sub-distal appendages of centrioles and ciliary membrane trafficking, in line with the proposed recent evolutionary origin of centrosomes and the intracellular pathway of ciliogenesis discussed above (see also Table S2B). A comparison of the present cluster with the original comparative genomics studies (Avidor-Reiss *et al*., 2004; Li *et al*., 2004) and more recent refined phylogenetic profiling approaches (Dey *et al*., 2015; Li *et al*., 2014; Nevers *et al*., 2017) revealed considerable overlap, but also components unique to one or other type of study (Fig. 2C, Table S2C). The enrichment of known ciliary components in the present study suggests our approach is similarly if not more effective at identifying *bona fide* components compared to other recent phylogenetic profiling approaches, although a direct comparison is complicated by the inclusion of known components as ‘seeds’ in the latter analyses.

**Figure 2:**
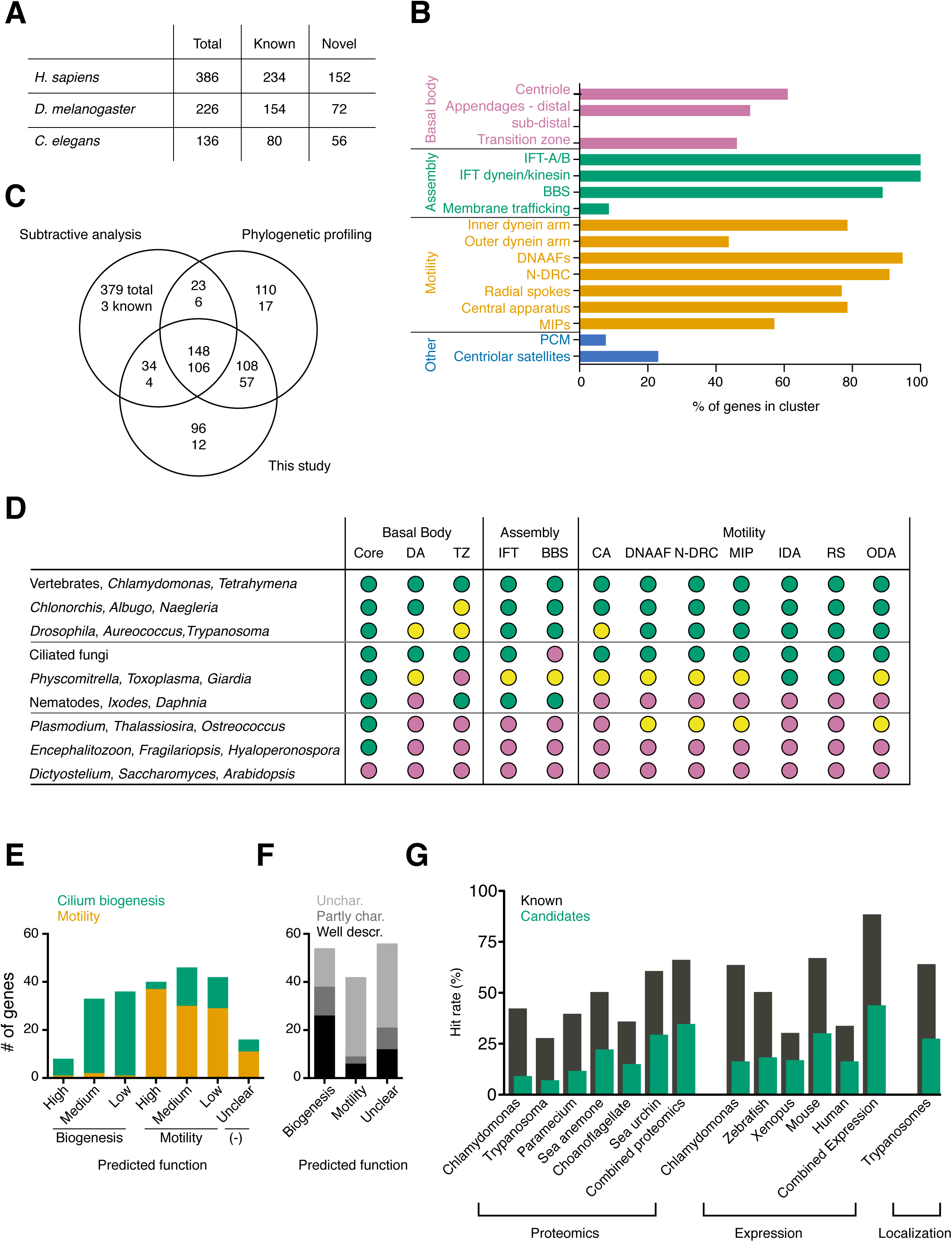
*In silico* analysis of the ciliary cluster. (**A**) Breakdown of ciliary cluster into known ciliary genes and those for which a ciliary function has yet to be established based on literature analysis. See also Table S2A. Not all 386 genes are conserved in *Drosophila* and *C. elegans* given the extensive gene loss in ecdysozoa (Guijarro-Clarke *et al*., 2020) and the loss of ciliary motility genes in nematodes. (**B**) Percentage of known genes for different functional categories associated with centrosomes and cilia represented within the ciliary cluster. See Table S2B for source data. (**C**) Venn diagram comparing the present 386 gene ciliary cluster with the output of previous comparative genomics analyses. In each case, the first number represent the total number of genes in each category, the second any centrosomal/ciliary gene found in the combined lists of genes defined in (B). Subtractive analysis refers to the original comparative analyses performed in 2004 (Avidor-Reiss *et al*., 2004; Li *et al*., 2004), combined into one dataset, phylogenetic profiling to the combined output of the more recent phylogenetic profiling studies (Dey *et al*., 2015; Li *et al*., 2014; Nevers *et al*., 2017). (**D**) Hierarchical clustering of species based on known genes within the ciliary cluster from the indicated functional categories shown in (B) reveals 9 distinct patterns of gene loss. Color code is green >2/3 of genes in indicated category present, yellow >1/3 of genes present, red <1/3 present. Representative species only shown in figure. See Table S2C for full list and source data. (**E**) Ciliary cluster genes can be sorted into predicted motility and biogenesis genes purely based on their pattern of conservation in species with motile and non-motile cilia (see Methods). Graph compares predicted function against actual, established function for all functionally characterized ciliary genes within the cluster (see annotation in Table S2A), grouped by level of conservation of the gene in ciliated species. The relatively high frequency of incorrectly assigned biogenesis genes at lower levels of conservation reflects their loss in ecdysozoa unrelated to their specific ciliary function. No prediction could be made for those genes present in <50% of ciliated species. See Table S2D for full details. (**F**) Predicted function of genes currently not linked to cilia based on phylogenetic footprint as in (E). Shading reflects overall level of functional characterization unrelated to any potential ciliary function. (**G**) Graph showing percent of known and candidate genes within the cluster found in selected large-scale proteomic, expression and localization studies. See Table S2E for full details.

Of the 386 genes in the ciliary cluster, all but one can be found in two or more of the 6 major eukaryotic groups other than unikonts shown in Fig. 1D (counting cryptophytes and haptophytes and disicristates and excavates each as one group). All genes therefore likely originate in the ancestral eukaryote and may have formed part of the original complement to assemble motile cilia. As discussed by David Mitchell in his 2016 review (Mitchell, 2016), diversification into modern organisms has involved not only acquisition of new features but also loss of existing ones, including part or all of the machinery to form cilia. Within the species represented in our analysis, cilia can be inferred to have been lost 13 times independently of each other over the course of evolution (4x in opisthokonts, 1x in amoebozoa, 3x in archaeplastidae, 2x in alveolates, 3x in stramenopiles, see Table S1A), which helps explain the clear definition of the ciliary cluster in Fig. 1E. Clustering species based on the major ciliary subcomplexes within the cluster reveals 7 other patterns (Fig. 2D and Table S2C). The machinery for ciliary motility but not cilia themselves has been entirely lost twice (once in nematodes, once in the arthropod *Ixodes scapularis*) and once in part (in the crustacean *Daphnia pulex*). These and other losses (e.g. of BBSome components in ciliated fungi) give rise to the grouping of genes functioning together in ciliary subcomplexes seen in Fig. 1E and can be used to predict the function of known genes based purely on their pattern of conservation with a high degree of accuracy (Fig. 2E and Table S2D). For the 152 novel candidate genes within the cluster, ∼2/3 can be assigned as putative ciliary assembly or motility genes, while for the remaining 1/3 the phylogenetic footprint is insufficient to make a confident prediction (Fig. 2F and Table S2D). But is there any experimental evidence for a ciliary function for these genes? Indeed, there is. As summarized in Fig. 2G, candidate genes have been identified in large scale proteomic, expression and localization screens performed in different experimental models, albeit unsurprisingly (given that those selfsame screens were used to isolate and characterize ciliary genes) at a lower rate than known genes within the cluster (see also Table S2E).

### Functional analysis of cluster genes reveals signature ciliary phenotypes in *Drosophila* and *C. elegans*

Based on our *in silico* analysis we were confident that within this cluster we had a set of genes highly enriched in key players in cilium assembly and motility. To test whether candidate genes indeed have a cilium-related function, we performed a comprehensive characterization by tissue-specific RNAi in *Drosophila* and mutant analysis in *C. elegans*. *Drosophila* is an attractive experimental model for such questions in that it possesses multiple types of cilia, including motile sperm flagella that have been reported to form in an IFT-independent manner in the cytoplasm, as well as sensory cilia that form in the canonical, IFT-dependent manner in olfactory and mechanosensory neurons (Han et al., 2003; Jana et al., 2016). Candidate genes can therefore be easily classified into genes required for general and compartmentalized cilium biogenesis and cilium motility. Using the GAL4-UAS system to deplete each candidate gene in the male germline (scoring male fertility), sensory neurons (assessing coordination in the negative geotaxis assay) and the whole animal (testing for non-specific, pleiotropic phenotypes), we compared the resultant phenotypes to those for known ciliary genes within the cluster (Fig. 3A-C). As can be seen in Fig. 3B, centriolar and ciliary transition zone components presented moderately strong phenotypes in both tissues, consistent with their role in both canonical and compartmentalized cilium biogenesis (Basiri et al., 2014; Jana et al., 2018; Vieillard et al., 2016). In contrast, IFT components presented strong phenotypes almost exclusively in neurons, confirming and generalizing their reported role specifically in compartmentalized cilium biogenesis (Han *et al*., 2003). This was also true for the IFT-associated BBSome, not previously functionally characterized in the fly, although phenotypes here were generally weak. In contrast, depletion of axonemal dyneins with the exception of certain outer arm dyneins strongly affected male fertility. Gravitaxis was also frequently perturbed, reflecting the role of dynein activity in mechanosensory neurons (Zur Lage et al., 2019). In total 88% of known ciliary genes within the cluster presented significant phenotypes in neurons, sperm or both (Fig. 3C, D, see also Table S3A). Remarkably, this was true also for 54 of 69 novel genes (78%). Depleting candidate genes in the whole animal tended to reveal no additional phenotypes, ruling out broader cellular functions. Given the limitations of the assays employed in this primary screen we conclude that most if not all novel candidate genes are indeed cilium-linked.

**Figure 3:**
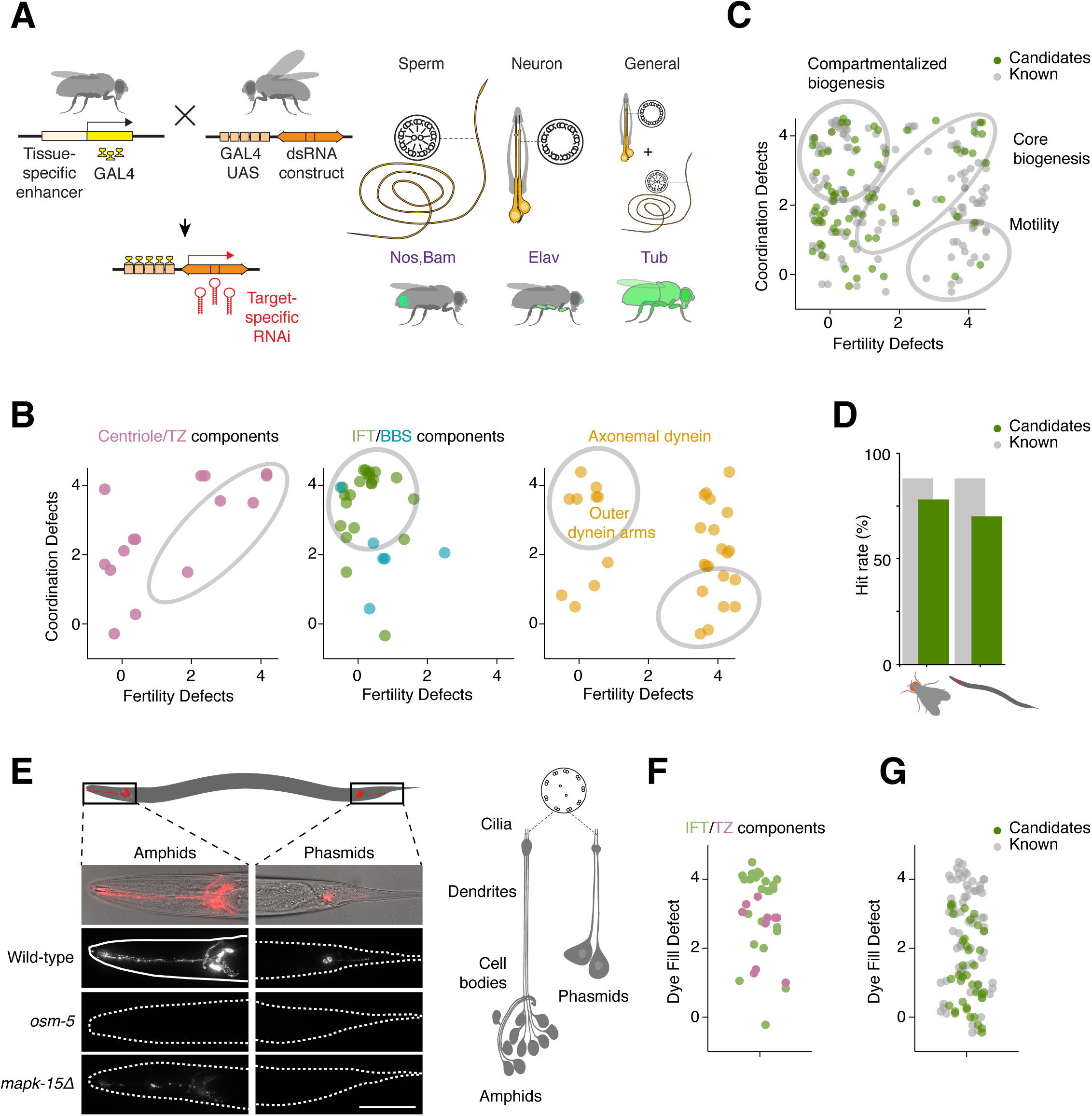
Results of primary screen in *C. elegans* and *Drosophila*. (**A**) In *Drosophila*, all conserved ciliary cluster genes were screened using the Gal4-UAS system for tissue-specific depletion by RNAi. Multiple RNAi lines per gene were crossed with drivers for neurons (Elav) and germline (Nanos and Bam combined), as well as a general driver (Tubulin). Flies were assayed for lethality, uncoordination and male fertility. Each phenotype was scored on a scale of 0 (no phenotype) to 4 (strong). See Table S3A for definitions. (**B**) Depletion of known centriolar and ciliary transition zone components results in defects in both coordination and male fertility, consistent with their core biogenesis function. In contrast, depletion of BBS and IFT components results in defects in neuronal function only, consistent with their role exclusively in compartmentalized biogenesis. Finally, depletion of axonemal dynein components primarily affects male fertility, although some outer arm dyneins also play significant roles in neuronal function. Each point on scatter plots represents one gene, with values shifted slightly (max +/-0.5) on both axes to avoid overlap of genes scoring identically. (**C**) Scatter plot displaying the phenotypes of all known (grey) and previously uncharacterised genes (green) within the ciliary cluster, plotted as in (B). See Table S3A for full details. (**D**) Graphical summary of percentage of known (grey) and novel genes (green) within the ciliary cluster presenting significant (i.e. with a score >1) ciliary phenotypes in *Drosophila* and *C. elegans*. Known and candidate genes scored similarly in both experimental models, suggesting a ciliary function for most if not all novel genes. (**E**) In *C. elegans*, all conserved ciliary cluster genes with available mutants as well as genes of particular interest for which mutants were generated in the course of this study were screened using the dye-fill assay to test for cilium structural integrity. 12 amphid neurons in the head and 4 phasmid neurons in the tail, both featuring non-motile sensory cilia, stain with the lipophilic dye DiI. Failure to take up dye is an indicator of ciliary structural defects. IFT mutants such as *osm-5(p813)* tend to display dye-fill null phenotypes, while other ciliary mutants present weaker phenotypes. Phenotypes were scored using a scale from 0 (no dye-fill phenotype) to 4 (null). See Table S3A for definitions. (**F**) Scatter plot displaying the phenotypes of mutants in IFT, BBS and transition zone components. Each point on scatter plots represents one gene, with values shifted slightly to avoid overlap. (**G**) Scatter plot displaying the phenotypes for all known (grey) and previously uncharacterised genes (green) within the ciliary cluster, plotted as in (F). See Table S3A for full details.

To confirm and extend our findings we turned to *C. elegans* as a complementary experimental model. Unlike *Drosophila*, *C. elegans* possesses exclusively non-motile cilia, present in post-mitotic sensory neurons, where they mediate the perception of chemosensory and mechanosensory stimuli. Compromised ciliary function leads to defects in complex behaviors such as chemotaxis, foraging and male mating. However, critically, cilia are dispensable for viability and fertility, allowing mutants to be propagated in their homozygous state (Inglis et al., 2007). To investigate the potential role of candidate genes in cilium assembly, we took advantage of the dye-fill assay as a simple and robust method to assess cilium structural integrity (Hedgecock et al., 1985). This assay is based on incubating worms with dilute solutions of the lipophilic dye DiI. In wild-type, this dye is taken up by a subset of neurons (12 amphid neurons in the head, 4 phasmid neurons in the tail) through their exposed ciliary endings and accumulates in their cell bodies (Fig. 3E). Defects in cilium assembly result in impaired dye-filling, which can be easily scored under the fluorescence microscope. For known ciliary genes within the cluster, the dye-fill assay revealed significant defects for 60 of 68 genes (88%), aided by the low animal to animal variability in *C. elegans*, with IFT mutants presenting stronger phenotypes than those affecting the transition zone (centriolar genes cannot be assayed given their essential role in embryonic development) (Fig. 3F, G). Remarkably, 26 of 37 candidate genes (70%) likewise presented a dye-fill phenotype. Phenotypes were generally weak, reflecting the fact that stronger, dye-fill null mutants have been previously identified in exhaustive large-scale screens using this assay (Perkins *et al*., 1986). With ecdysozoa and particularly nematodes having experienced widespread gene-loss (Guijarro-Clarke et al., 2020), including within the ciliary cluster, not all 386 genes could be assayed in *Drosophila* or *C. elegans*. Our results are, however, consistent with our hypothesis that essentially all cluster genes function in some aspect of cilium assembly or motility.

### Secondary analysis identifies candidate genes involved in the early stages of ciliogenesis

Although our primary screen in *Drosophila* allowed the identification and general classification of candidate genes, the precise nature of the process disrupted in each case remained to be determined. Thus, defects in male fertility could arise from a lack of sperm flagella or lack of sperm motility. Similarly, neuronal defects could arise from a lack of cilia or lack of mechanosensory function. To classify novel ciliary genes according to their function and identify candidates for further study we employed a set of secondary assays for both sperm and neuronal cilia, comparing their depletion phenotypes to those for representatives of the various functional classes. For any gene scoring significantly in the coordination assays as well as those chosen for further characterization based on their spermatogenesis phenotype, the chordotonal organs responsible for proprioception located in the animals’ legs (Fig. 4A, B) were dissected and examined by DIC microscopy, as well as by immunofluorescence microscopy using antibodies to markers for the apical ciliary membrane (NompC) and basal bodies (Sas-4) (Dobbelaere *et al*., 2020). Chordotonal organs are made up of multiple scolopidia, each of which contains a pair of ciliated nerve endings ensheathed by a glial cell and attached with their ciliary tips to the cuticle via a cap cell (Kernan, 2007). We found defects in cilium biogenesis to manifest themselves in different ways, with depletion of centriole biogenesis and IFT components resulting in the total absence or reduced numbers of cilia, while depletion of other components yielded more subtle defects including shorter (as assessed by the position of the ciliary dilation) or morphologically abnormal cilia (Fig. 4B). Depletion of candidate genes primarily resulted in the latter type of defect, with 8 of 36 genes examined displaying shortened or abnormal cilia (Selected examples shown in Fig. 4B, C, quantitation in Table S3B). Supernumerary cilia, formed by conversion of daughter centrioles into mothers, a phenotype associated with depletion of Centrobin (Gottardo *et al*., 2015), were not observed for any candidate gene.

**Figure 4:**
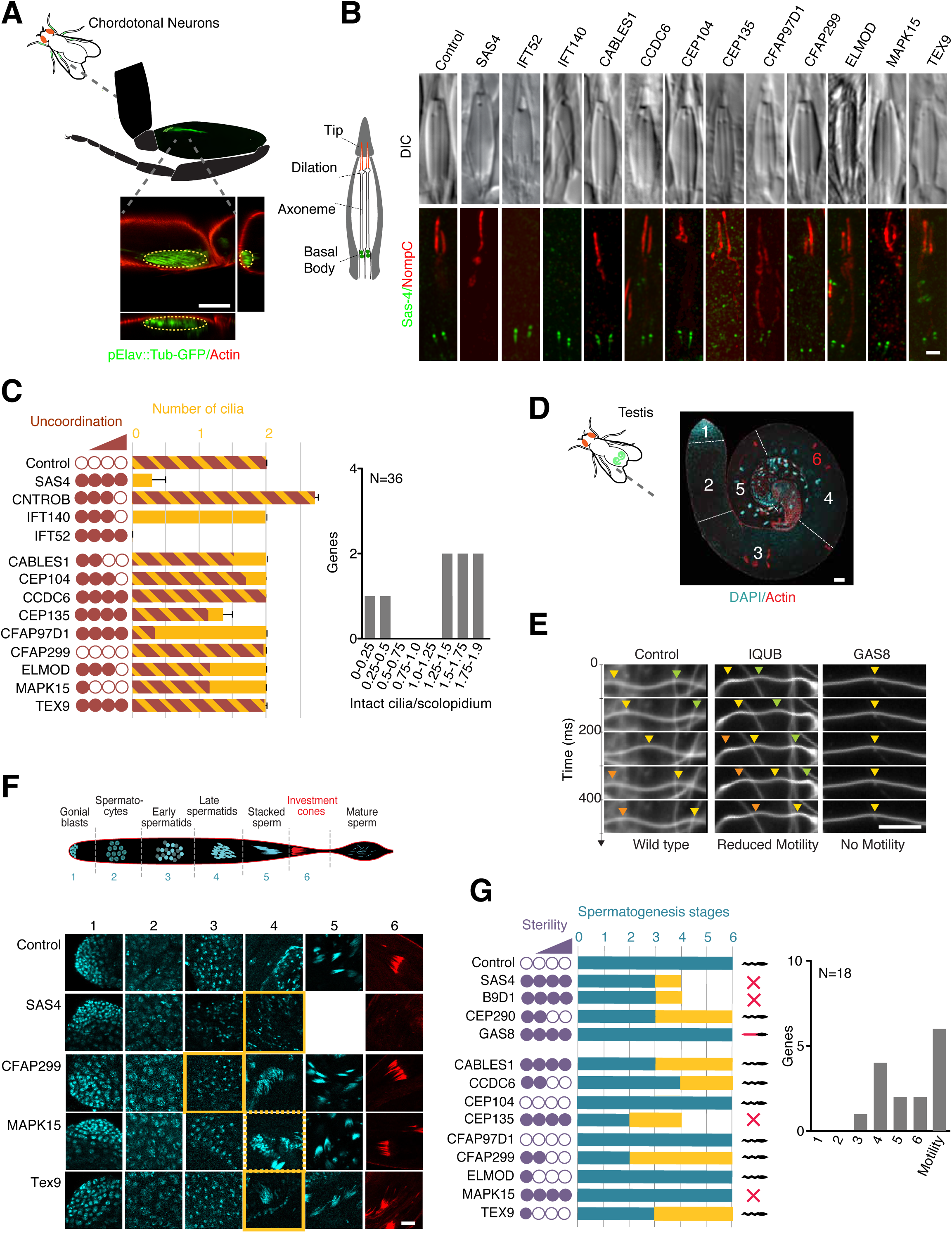
Results of secondary screen in *Drosophila*. (**A**) Schematic and confocal images of leg chordotonal neurons, visualized using pElav-Gal4;UAS-Tubulin-GFP and actin staining, with xy, xz and yz image projections to show position relative to the cuticle of the leg. (**B**) Schematic and confocal images of scolopidia in chordotonal organ of the fly. Each scolopidium contains two cilia with their ends embedded in a cap cell. Scolopidium outline, cilia and ciliary dilation can be visualized by DIC. Immunofluorescence micrographs with Sas4 and NompC antibodies used to mark basal bodies and ciliary tips, respectively. DIC images of flies depleted of candidate genes by RNAi reveals relatively few defects on par with IFT genes such as IFT140, with the exception of misplaced dilations for CFAP97D1 and Cables1. NompC/Sas-4 staining reveals more subtle phenotypes, including mislocalized NompC signal (with CFAP97D1, Cep135, Elmod and Mapk15) and apparent ciliary structural defects (Cep104 and Cables1). (**C**) Graph summarizing neuronal defects observed for candidate genes chosen for further analysis, with selected known centriolar/ciliary genes shown for comparison. Yellow bar is based on DIC analysis, brown bar based on NompC/Sas-4. Uncoordination score is from primary screen. The second graph plots the number and severity of neuronal ciliary defects (DIC and NompC/Sas-4 combined) for all previously uncharacterized ciliary genes chosen for further analysis. (**D**) Image of dissected control testis stained for DNA and actin, showing the spatial separation of the different stages of spermatogenesis from stem cells (1) to bundles of mature sperm (5), each with their own distinct nuclear morphology. Also visible are the actin cones that strip away extra cytoplasm during the later stages of spermatogenesis. (**E**) Mature sperm can be dissected from the seminal vesicle and motility analyzed using high speed video capture in dark field microscopy. Sinusoidal motion can be seen in wild type. In RNAi depletions of certain known motility genes (here IQUB and GAS8), this motion is reduced or even absent. (**F**) Spermatogenesis in wild-type and RNAi depletions of selected genes, highlighting the stage where morphogenesis first becomes noticeably abnormal. (**G**) Graph summarizing sperm defects observed for candidate genes chosen for further analysis, with selected known centriolar/ciliary genes shown for comparison. Yellow bar indicates stage where phenotypes first become apparent, blue bar indicates furthest progression observed. Any sperm accumulating in the seminal vesicle were scored for motility. The second graph plots the severity of sperm ciliary defects (stage at which defects were first observed) for all previously uncharacterized ciliary genes chosen for further analysis. Scalebars are 50µm (A, D), 1µm (B), 20µm (E, F). Error bars in (C, G) are standard deviation.

For any gene scoring significantly in the male fertility assay as well as those chosen for further characterization based on their neuronal phenotype, the process of spermatogenesis was examined in two ways. First, testes were dissected, fixed and stained to visualize DNA and the actin cytoskeleton. This allowed us to monitor the different stages of sperm differentiation using nuclear morphology, as well as the formation of actin cones during individualization (Fig. 4D). Second, we dissected the seminal vesicle where mature sperm are stored and filmed the movement of sperm tails using dark field microscopy (Fig. 4E). Examining known ciliary genes, we found that centriolar and transition zone components gave the strongest phenotypes, arresting sperm morphogenesis at early stages of differentiation, whereas motility genes only affected later stages of differentiation (Fig. 4F, G). Interestingly, sperm found in the seminal vesicle was generally found to be at least partly motile, with defects in ciliary motility primarily manifesting themselves in an empty seminal vesicle as sperm fail to move from the testes to this storage organ. Depletion of candidate genes resulted in a wide spectrum of phenotypes, with some genes resulting in defects as early as the early spermatid stage, similar to Sas4 and CEP290, while others progressed as far as the individualization stage. Overall, 15 of 18 genes examined yielded significant phenotypes in this assay (Fig. 4F, G, quantitation in Table S3C).

In total, 19 of 37 genes (51%) examined based on their primary screen phenotype presented clear defects in either neurons or sperm. In our further characterization we focused on a handful of those genes which presented particularly strong phenotypes consistent with a function in early stages of ciliogenesis and/or displayed an evolutionary conservation pattern consistent with a central role in cilium biogenesis.

### CABLES1, CCDC6 and TEX9 as novel basal body components related to Bld10/CEP135

Amongst the candidate genes in the screen that caught our attention were CABLES1, CCDC6 and TEX9, which presented significant phenotypes in neurons as well as sperm in our primary and secondary analyses. TEX9 has previously been localized to centriolar satellites and the ciliary base in vertebrate cells (Gupta et al., 2015; Nevers *et al*., 2017), but remains functionally uncharacterized. CCDC6 has an extensive literature as a recurrent fusion partner for the oncogenic tyrosine kinases ROS1 and RET (Cerrato et al., 2018), yet itself remains entirely uncharacterized, although interestingly it was reported amongst the interactors in pulldowns of the IFT-B component Cluap1/IFT38 (Beyer et al., 2018). Finally, CABLES1 has been reported to interact with CDK5 and c-ABL and linked to control of cell proliferation and/or differentiation (Wang et al., 2010; Zukerberg et al., 2000), with no apparent connection to centrosomes or cilia. Given the severity of the neuronal and spermatogenesis phenotypes, ultrastructural analysis was performed on RNAi-depleted animals (and in the case of CABLES1 also deletion mutants) for all three candidate genes. In the case of neuronal cilia, we observed a general disorganization of ciliary tips, with missing axonemal doublet microtubules (Fig. 5A). More striking was the phenotype in sperm, where the central pair of microtubules was frequently missing, while the outer microtubule doublets were unaffected (Fig. 5B). Such a phenotype has hitherto been reported in *Drosophila* only for a single gene, the centriolar cartwheel component Bld10/CEP135 (Carvalho-Santos et al., 2012). A re-examination of CEP135 by RNAi and mutant analysis confirmed these earlier observations but also revealed defects in mechanosensation and ciliary ultrastructure in chordotonal neurons, similar to CABLES1 and CCDC6 (Fig. 5A). We should note that the latter finding stands in contrast to initial reports on this gene in the fly which reported no apparent defects in coordination (Mottier-Pavie and Megraw, 2009), which we attribute to the use of a stronger loss of function mutant than in the original work.

**Figure 5:**
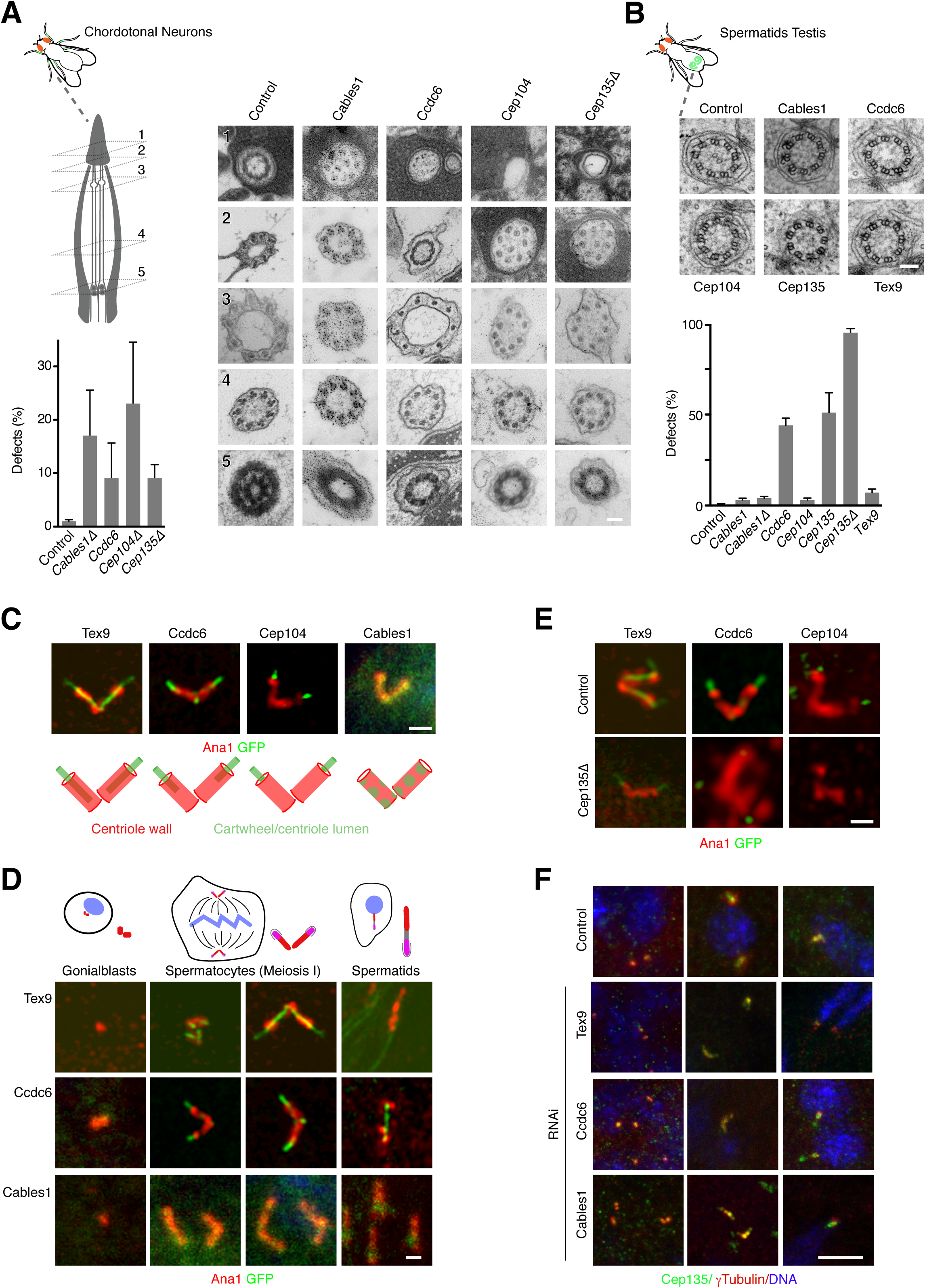
CABLES1, CCDC6 and TEX9 as novel basal body components related to Bld10/CEP135. (**A**, **B**) RNAi-mediated depletion/deletion of CABLES1, CCDC6, CEP104 and TEX9 in *Drosophila* chordotonal neurons (A) and sperm (B) results in defects in ciliary ultrastructure similar to CEP135. In neurons, there is a loss of 9-fold symmetry towards the ciliary tips and axonemes fail to reach the cap cell. In sperm, there is a loss of the inner pair of microtubules, a phenotype hitherto only associated with CEP135. (**C**) GFP fusions to TEX9, CCDC6 and CEP104 localize to the central lumen of sperm basal bodies marked by Ana1, while CABLES1 only faintly localizes to basal bodies. (**D**) TEX9, CCDC6 and CABLES1 are recruited to maturing basal bodies during *Drosophila* spermatogenesis. TEX9 and CCDC6 images in panel 3 from (C). (**E**, **F**) Localization interdependencies between CEP135, CEP104 and the novel components TEX9, CCDC6 and CABLES1 in spermatogenesis. CEP135 is required for the localization of TEX9, CCDC6 and CEP104 to mature basal bodies (E), while CEP135 localization is independent of TEX9, CCDC6 and CABLES1 at all stages of spermatogenesis (F). Scalebars are 100nm (A, B), 1µm (C-E), 5µm (F). Error bars in (A, B) are standard deviation.

This phenotypic similarity to CEP135 prompted us to examine the localization of our novel components in the fly, with GFP transgenic fusions expressed under the general Ubq promoter. Remarkably, GFP fusions for both TEX9 and CCDC6, like CEP135 (Tian et al., 2021), localized to the center of the centriole barrel marked by Ana1, but extending further along the length of the extended spermatocyte basal body (Fig. 5C). The third protein, CABLES1, also localized to spermatocyte basal bodies, albeit much more weakly, precluding more detailed analysis. No localization was observed to centrosome-organizing centrioles elsewhere in the fly. Instead, all three proteins were recruited as centrioles convert into basal bodies during spermatogenesis (Fig. 5D). In the course of our studies we came across one other known ciliary cluster gene, CEP104, which shared some of the phenotypic features of the above-named components. Ultrastructural analysis of both RNAi-depleted and mutant animals revealed the same signature ciliary defects in sperm and neurons (Fig. 5A, B). CEP104 in vertebrates and *Chlamydomonas* has been shown to be a microtubule plus-end tracking protein that translocates during ciliogenesis from the distal end of centrioles to the ciliary tip (Jiang et al., 2012; Satish Tammana et al., 2013). Cep104 in *Drosophila* shares this localization, forming a central focus at the distal end of sperm basal bodies that co-localizes with the most distal populations of CCDC6 and TEX9 (Fig. 5C).

We then have four (with CABLES1 potentially five) components that share a localization to the core of basal bodies, with CEP135 at the proximal end and CEP104 at the distal end, and function in ciliogenesis, particularly in formation of the central pair and organization of ciliary tips. We therefore propose that these components form part of a common ciliogenesis pathway with CEP135 at its base. Consistent with such a pathway, loss of CEP135 is independent of CABLES1, CCDC6 and TEX9 (Fig. 5F), but strongly affects the localization of all other components without affecting centriole assembly *per se* (Fig. 5E).

### CFAP97D1 as part of a family of CFAP97 domain-containing proteins required for axoneme assembly/stability

CG14551/CFAP97D1 drew our attention due to its strong phenotype specifically in neurons. CG14551 is the only *Drosophila* ortholog of vertebrate CFAP97D1 and 2, part of a family of proteins that also includes CFAP97/KIAA1430 (Hemingway in *Drosophila*) (Fig. 6A). CFAP97D1 in mice has been found to be required for sperm flagellar axoneme integrity and male fertility (Oura et al., 2020), while its paralog CFAP97D2 remains uncharacterized. Similarly, Hemingway was found be required for sperm flagellum assembly and ciliary motility in the fly, including in auditory sensory neurons, but with no apparent defects in coordination (Soulavie et al., 2014) (vertebrate CFAP97 is presently uncharacterized). It was then surprising to see a phenotype for CFAP97D1 exclusively in neurons. Further characterization including by mutant analysis confirmed this specificity. Ultrastructural analysis of chordotonal cilia revealed broken axonemes and missing axonemal microtubule doublets, particularly towards the ciliary tip (Fig. 6B), a phenotype reminiscent to what has been observed for sperm flagella with vertebrate CFAP97D1 and *Drosophila* Hemingway (Oura *et al*., 2020; Soulavie *et al*., 2014). CFAP97D1 was not localized in that study, while staining for Hemingway revealed no apparent localization to sperm. In contrast, *Drosophila* CFAP97D1 showed a clear ciliary localization, particularly in the area of the ciliary dilation towards the tip of chordotonal cilia (Fig. 6C). Such a localization is consistent with the proposed function of CFAP97 domain-containing proteins in cilium elongation/stability. Our data suggests that this function is not specific to motile cilia/flagella. Consistent with this, the Human Protein Atlas (proteinatlas.org, (Uhlén et al., 2015)) notes that while CFAP97D1 is testis specific, CFAP97D2 is more broadly expressed in ciliated tissues, including those bearing exclusively non-motile cilia such as the pancreas, while CFAP97 is essentially ubiquitous.

**Figure 6:**
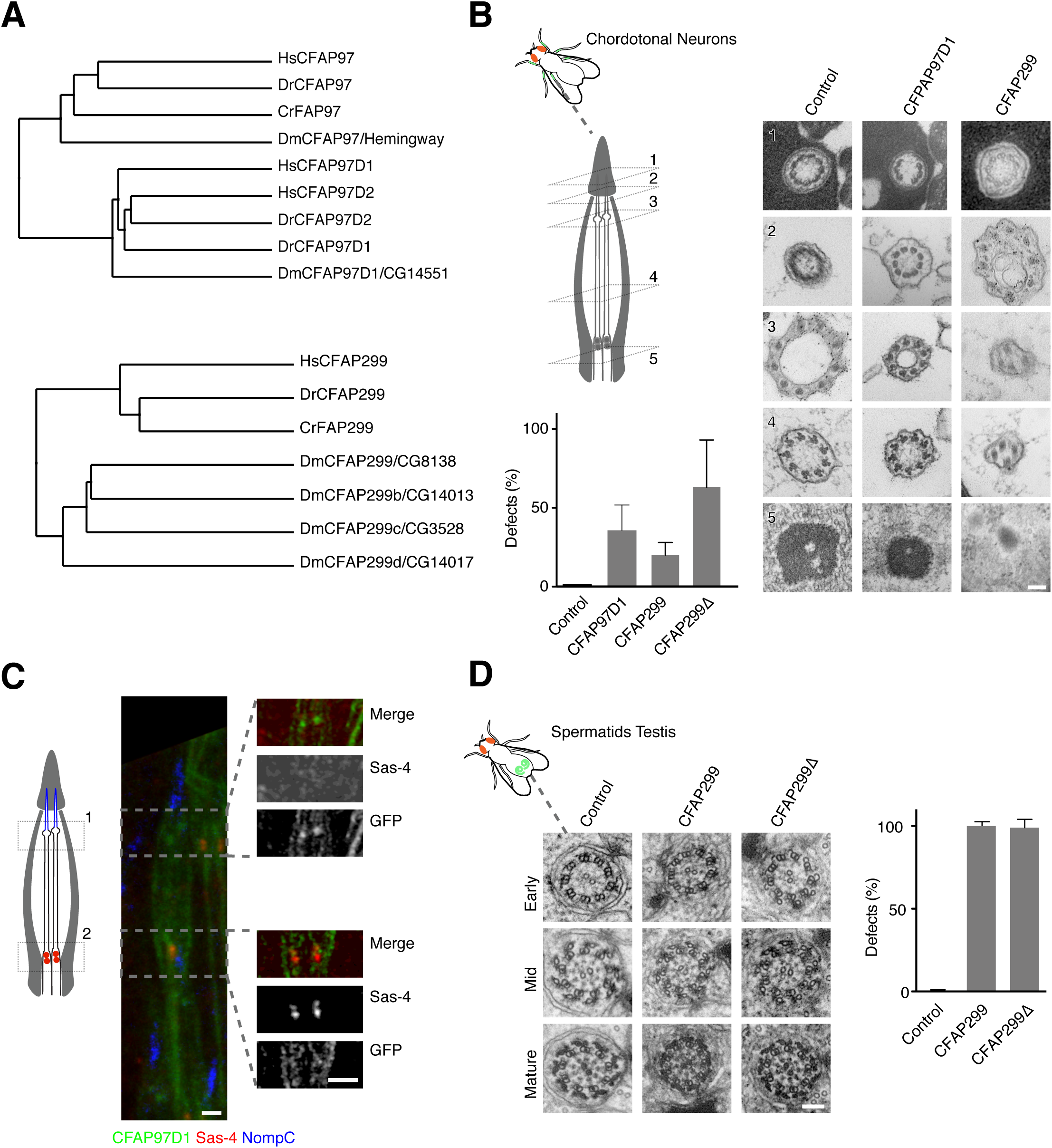
A conserved role for CFAP97 and CFAP299 family members in axoneme assembly and ciliogenesis. (**A**) Relationship between CFAP97 and CFAP299 family members in humans (Hs), zebrafish (Dr), *Chlamydomonas* (Cr) and *Drosophila* (Dm). Average distance trees calculated using the BLOSUM62 substitution matrix. *Drosophila* CFAP97D1 and its vertebrate paralogs CFAP97D1 and CFAP97D2 are distantly related to CFAP97/Hemingway, while CFAP299 has 4 paralogs in *Drosophila* with distinct tissue expression patterns. Characterized here is the most closely related ortholog, CG8138, hereafter CFAP299. Accession numbers: HsCFAP97 (NP_065878.1); DrCFAP97 (NP_997943.1); CrFAP97 (XP_042919651.1); DmCFAP97/Hemingway (NP_650714.1); HsCFAP97D1 (NP_001129955.1); HsCFAP97D2 (XP_016876399.1); DrCFAP97D1 (XP_697280.2); DrCFAP97D2 (XP_005167838.1); DmCFAP97D1/CG14551 (NP_651462.1); HsCFAP299 (XP_047305931.1); DrCFAP299 (NP_001108596.1); CrFAP299 (XP_001697404.2); DmCFAP299/CG8138 (NP_650260.1); DmCFAP299b/CG14013 (NP_608941.1); DmCFAP299c/DmCG3528 (NP_608686.1); DmCFAP299d/DmCG14017 (NP_608942.1). (**B**). RNAi-mediated depletion/deletion of *Drosophila* CFAP97D1 and CFAP299 results in defects in ciliary ultrastructure in chordotonal neurons, including broken axonemes and perturbed 9-fold symmetry. At least 3 animals examined per condition. (**C**) A GFP fusion to CFAP97D1 localizes to the ciliary dilation but not basal body in chordotonal neurons. Insets acquired in Airyscan mode. (**D**) RNAi-mediated depletion/deletion of CFAP299 results in failure of membrane deposition around the developing sperm axoneme. At least 3 animals examined per condition. Scalebars are 100nm (B, D), 1µm (C). Error bars in (B, D) are standard deviation.

### Unique role for CFAP299 in ciliary membrane deposition and ciliogenesis

CG8138/CFAP299 along with CG14013, CG14017 and CG3528 is one of four *Drosophila* orthologs of vertebrate C4orf22/CFAP299 (Fig. 6A), a largely uncharacterized protein reported to be testis-enriched and tentatively linked to spermatogenesis (Li et al., 2019a), but according to the Human Protein Atlas (proteinatlas.org, (Uhlén *et al*., 2015)) also expressed in other ciliated tissues including the lung and oviduct. CG8138 initially presented with a strong phenotype in male fertility in the primary and secondary screen. Ultrastructural analysis of both RNAi-depleted and mutant animals revealed what to our knowledge is a unique and unprecedented phenotype: a seemingly intact flagellar axoneme but a total failure of membrane deposition to complete sperm individualization (Fig. 6C). As described above, *Drosophila* spermatogenesis is highly unusual in that flagella form in an IFT-independent manner in the cytoplasm and are then extruded in a process reminiscent of what is believed to occur in the malarial parasite *Plasmodium* (Han *et al*., 2003; Sinden et al., 1976). The proteins responsible for axoneme elongation in the cytoplasm, whether unique to that context or shared with canonical compartmentalized ciliogenesis, are not known. Tantalizingly, CFAP299 is also conserved in *Plasmodium*, yet its wider phylogenetic distribution, including in humans, suggest a function also in canonical ciliogenesis. Indeed, CG8138/CFAP299 mutant animals also presented severe defects in chordotonal neurons, with highly aberrant axonemes lacking nine-fold symmetry (Fig. 6B). CG8138 shares extensive homology with its paralogs CG14013, CG14017 and CG3528, also putatively expressed in neurons and sperm, raising the possibility of functional redundancy. Consistent with this, weaker phenotypes were also observed for two other more distantly related paralogs CG14013 and CG3528 (see Table S3A). How CFAP299 and its orthologs contribute to ciliogenesis is as yet unclear. No specific localization was observed for N- and C-terminal GFP fusions in either sperm or neurons. However, its presence in the *Chlamydomonas* flagellar proteome (Pazour *et al*., 2005) suggests a role directly in cilium extension.

### MAPK15 and ELMOD as highly conserved regulators of ciliogenesis

The final two genes, MAPK15 and ELMOD, have to different degrees been previously characterized, but were nevertheless chosen for further analysis based on the type of protein they encode, their highly conserved nature and the severity of the phenotype associated with their depletion/mutation. MAPK15 (also known as ERK7 or 8) is an atypical mitogen-activated protein kinase, atypical in the sense that it does not function as part of a classical three-tiered MAPKKK–MAPKK–MAP kinase cascade (Coulombe and Meloche, 2007). Originally linked to cellular homeostasis and the maintenance of genomic integrity (Deniz et al., 2022), MAPK15 was more recently found to also regulate ciliogenesis by promoting the apical migration of basal bodies in *Xenopus*, reportedly by phosphorylating the actin regulator CapZIP (Miyatake et al., 2015). In *C. elegans*, putative *mapk-15* loss of function mutations were found to perturb neuronal morphogenesis (McLachlan et al., 2018) but also ciliary architecture as well as the localization of many ciliary proteins (Kazatskaya et al., 2017; Piasecki et al., 2017). Ciliogenesis was also reported to be affected in human RPE1 cells (Kazatskaya *et al*., 2017). With the kinase localizing to the ciliary base, MAPK-15 was proposed to act in a gating capacity to regulate ciliary trafficking and thereby ciliogenesis (Kazatskaya *et al*., 2017).

What caught our attention is that MAPK-15 is one of only 14 genes universally conserved in ciliated species and lost in non-ciliated ones, a remarkably short list that also includes the core centriolar structural components SASS6 and CENPJ/SAS-4, suggesting a central role in cilium biogenesis (see Table S2D). Consistent with this, MAPK15 scored strongly in the primary and secondary screens in the fly as well as in the worm. Given that available mutants in *C. elegans* are only partial gene deletions, a complete knockout was generated by CRISPR/Cas9-mediated gene editing. This mutant presented an even stronger phenotype (with an average of 4.3 and 0.3 dye-filled neurons in amphids and phasmids, respectively, compared to 6.9 and 0.4 for the original mutant), with cilium length and dendrite extension (indicative of transition zone dysfunction in the worm (Schouteden et al., 2015)) both strongly reduced (Fig. 7A, B). Ultrastructural analysis revealed that *mapk-15* null mutants display severe defects in transition zone organization and axoneme elongation, with additional distortions of the periciliary membrane previously noted for loss of the microtubule organizing center at the ciliary base (Garbrecht et al., 2021) (Fig 7C-E). This combination of phenotypes in the worm is remarkable in that IFT mutants do not affect transition zone organization, while loss of transition structures barely affects cilium elongation (Schouteden *et al*., 2015) (Fig. 7A, B). Indeed, no other previously characterized *C. elegans* gene shares this dual phenotype. The *Drosophila* phenotype is likewise remarkable: severe defects in both sperm and neuronal axoneme elongation and structural integrity (Fig. S2A, B). Defects in flagellar morphogenesis in particular are telling, given that this process is IFT-independent in the fly. MAPK15 thus clearly functions in IFT-independent step in axoneme elongation, consistent with its conservation in *Plasmodium*, which entirely lacks IFT components. MAPK15 has previously been reported to localize to the (acentriolar) ciliary base in *C. elegans* and basal bodies in vertebrates, as well as to the periciliary membrane compartment in *C. elegans* (Kazatskaya *et al*., 2017). We could confirm those observations for the worm (Fig. 7F, S1A) and extend them to *Drosophila*, where MAPK15 co-localized with the core centriolar component Sas4 to basal bodies in both chordotonal neurons and sperm (Fig. S2C, D). What is worth noting, however, is that MAPK15 does not localize to centrioles elsewhere in the worm or fly. Instead, MAPK15 is recruited to basal bodies at a time comparable to the expression of the very first ciliary components (1.5-fold stage in *C. elegans* embryogenesis (Serwas *et al*., 2017) (Fig. S1B), meiosis I in *Drosophila* spermatogenesis (Basiri *et al*., 2014) (Fig. S2C). In summary, then, what MAPK15 represents is a universally conserved regulator that is recruited at the onset of ciliogenesis to initiate the assembly of otherwise largely independent ciliary structures (the ciliary transition zone and axoneme).

**Figure 7:**
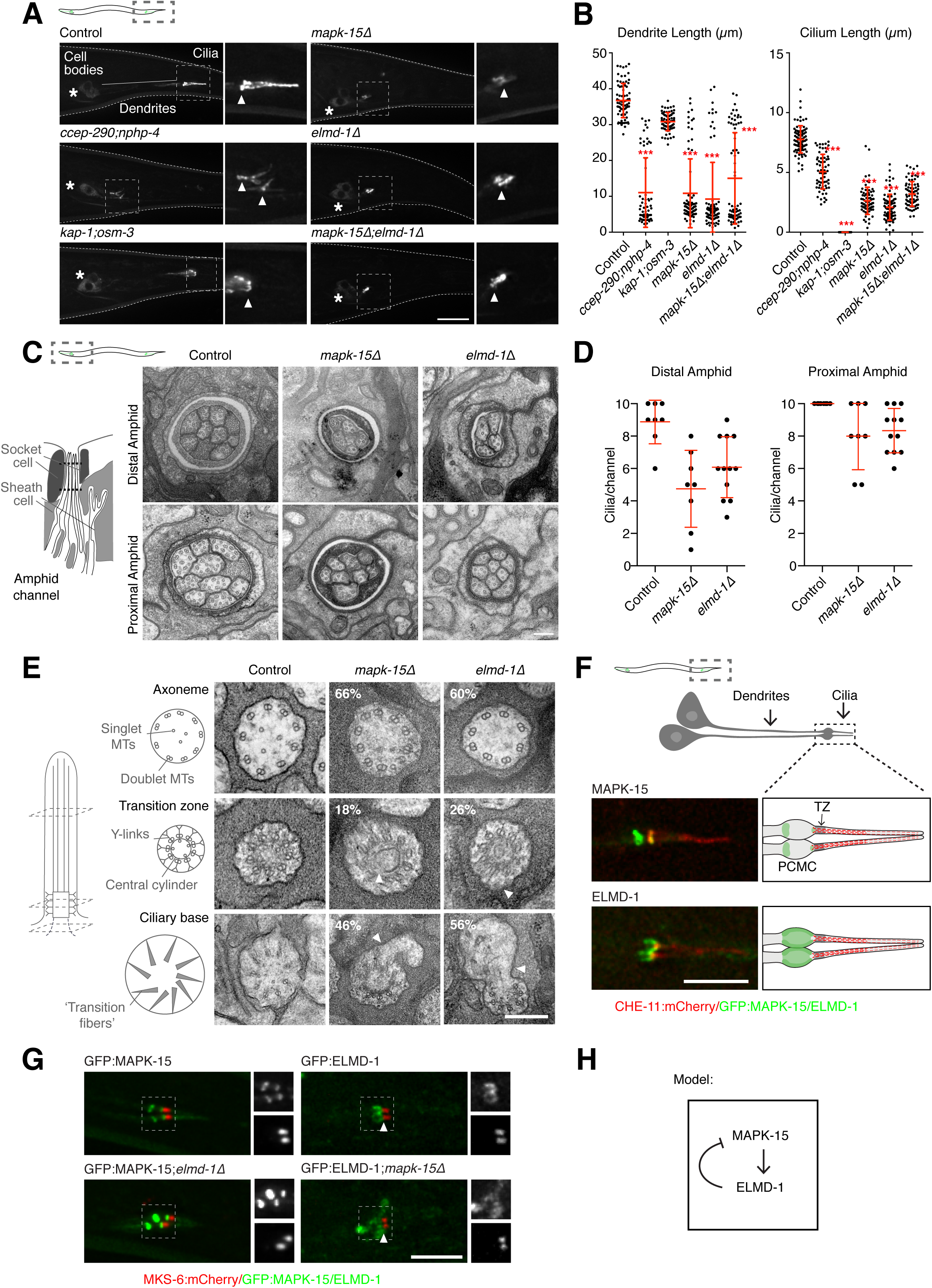
MAPK15 and ELMOD as highly conserved regulators of cilium biogenesis. (**A**, **B**) Full gene deletions of *mapk-15* and *elmd-1* in *C. elegans* result in both shortened cilia and shortened dendrites, combining the signature phenotypes of IFT and transition zone mutants (here *kap-1;osm-3* and *ccep-290;nphp-4* double mutants, respectively). Representative images of phasmid (tail) neurons expressing OSM-6:GFP (A) and quantitation of cilium and dendrite lengths (B) in control and mutant animals as indicated. n >20 animals for all conditions. Asterisks indicate statistically significant difference to wild-type (t-test, P < 0.001). *mapk-15;elmd-1* double mutants do not display phenotypes stronger than either single mutant (t-test, not significant). (**C**-**E**) *mapk-15* and *elmd-1* mutants display defects in ciliary ultrastructure. (C) Cross-sectional views through the amphid channel reveal defects in cilium extension in both mutants, with fewer cilia protruding into the channel. (D) Quantitation of cilium extension defects at the level of the proximal and distal amphid channel. (E) Higher magnification views of individual cilia reveal defects at the level of the proximal segment of the axoneme (fewer than 9 doublet microtubules), transition zone (disorganization of the central cylinder and overall architecture) and ciliary base (membrane blebs). Percentages reported in figure based on examination of 4 *mapk-15* and 7 *elmd-1* mutants. (**F**) Endogenous/Endogenous promoter GFP fusions to ELMD-1 and MAPK-15 localize to the ciliary base and periciliary membrane compartment in *C. elegans* phasmid neurons, the former marked by the IFT component CHE-11. (**G**, **H**) Localization interdependencies between MAPK-15 and ELMD-1. (G) Loss of ELMD-1 results in a hyperaccumulation of MAPK-15 at the *C. elegans* ciliary base, while loss of MAPK-15 impairs localization of ELMD-1, suggesting that (H) MAPK-15 functions upstream of ELMD-1. Scalebars are 10µm (A), 200nm (C, E), 100nm (E, F), 5µm (F, G), 1µm. Error bars in (B, D) are standard deviation.

ELMOD1-3, represented in *Drosophila* and *C. elegans* by a single ortholog, ELMOD/ELMD-1, are part of a family of ELMO (Engulfment and cell motility) domain-containing proteins, named after its founding member CED-12/ELMO1, which acts to promote apoptotic cell engulfment by activating CED-10/Rac1 GTPase (Wu et al., 2001; Zhou et al., 2001). In contrast, ELMOD family members are thought to act as ARF/ARL GAPs, with broad cellular effects, including on the actin cytoskeleton (Li et al., 2019b). All three vertebrate ELMOD proteins have recently been localized to basal bodies and linked to ciliogenesis, albeit with seemingly opposing effects: While loss of ELMOD1 and 3 reduced ciliation in mouse embryonic fibroblasts, loss of ELMOD2 actually resulted in an increase, with those cilia displaying abnormal morphology, including apparent splaying of axonemes and ciliary rootlet defects (Turn et al., 2022; Turn et al., 2021) (ultrastructural analysis has so far not been performed). Common to all three perturbations is a mislocalization of ciliary proteins, suggesting ciliary trafficking is at least partly responsible for the observed defects (Turn *et al*., 2022; Turn *et al*., 2021). In our primary and secondary screen in *Drosophila* ELMOD presented with strong phenotypes in both sperm and neurons and rather less of a phenotype in *C. elegans*. However, the latter result was based on a nonsense mutant of uncertain consequence (*elmd-1(gk386113)*, Trp24Ter, (Thompson et al., 2013)). Similarly weak phenotypes were reported recently for another mutant, *elmd-1(syb630)*, a partial gene deletion (Cevik et al., 2021). We therefore obtained a full gene deletion, which yielded a much stronger phenotype (with an average of 4.9 and 0.5 dye-filled neurons in amphids and phasmids, respectively, compared to 10.4 and 3.8 for the original mutant) and was used for further analysis. Remarkably, in both *C. elegans* and *Drosophila* ELMOD presented phenotypes nearly indistinguishable from MAPK15. Thus, in *C. elegans* both cilium and dendrite lengths were much reduced, with axonemal, transition zone and periciliary membrane compartment defects matching those of MAPK15 (Fig. 7A-E). Likewise, in *Drosophila* ELMOD depletion led to defects in axoneme elongation and structural integrity in both sperm and chordotonal neurons (Fig. S2A, B). ELMOD and MAPK15 localization was also very similar if not quite identical: In *Drosophila*, both proteins localize to basal bodies (Fig. S2C, D), while in *C. elegans* both display a dual localization to the ciliary base as well as the periciliary membrane compartment, although here the domain occupied by ELMOD is more extensive (Fig. 7F, S2A). Finally, neither protein localizes to centrioles, with both being recruited to basal bodies early in ciliogenesis with near-identical timing in *Drosophila* sperm and *C. elegans* neurons (Fig. S1B, S2C). These similarities led us to posit that these proteins may function in the same molecular pathway. Consistent with this, *elmd- 1;mapk-15* double mutants in *C. elegans* presented a phenotype no more severe than that of either single mutant (Fig. 7A, B). Finally, we sought to investigate whether there were any localization interdependencies between these two ciliary components. We found loss of MAPK-15 to impair proper ciliary targeting of ELMD-1, while MAPK-15 localization was not affected by loss of ELMD-1 (indeed, levels are markedly increased) (Fig. 7G). Our results therefore indicate that ELMD-1 functions downstream of MAPK-15 in a ciliogenesis program that serves to convert centrosome-organizing centrioles into basal bodies capable of supporting axoneme extension and transition zone assembly (Fig. 7H).

## DISCUSSION

Cilia in many ways are ideally suited for comparative genomics approaches in that they are evolutionarily ancient yet not universally conserved in all extant species. Once cilia are no longer present, gene loss follows very rapidly, as demonstrated by the example of *Emiliania huxleyi*, where haploid-specific genes including those required for cilium biogenesis have been found to have been lost multiple times in different sub-populations of the same species (von Dassow et al., 2015). The resultant presence and absence pattern for ciliary genes in the genomes of fully sequenced eukaryotes creates a prominent phylogenetic signature that sets them apart from other highly conserved genes. We found that, at least for cilia, a simple but robust phylogenetic profiling approach based on bidirectional best matches and hierarchical clustering outperforms more sophisticated approaches and identifies a single cluster of 386 genes. This cluster includes the vast majority of known players involved in key steps of centriole assembly, cilium biogenesis and motility, as well as a set of novel genes that our functional analysis indicates are likewise associated with different aspects of cilium assembly and motility. We believe this set to constitute the core complement of genes required to assemble motile cilia. The phylogenetic distribution of these 386 genes indicates that all of them originate in the last eukaryotic common ancestor. In contrast to the rather simple molecular architecture of centrioles (Pelletier et al., 2006), the larger basal body-cilium superstructure of which they form an integral part then is highly complex and has been for >1000 million years.

Importantly, we have carried out a comprehensive verification of our bioinformatic predictions by performing a systematic RNAi-based analysis in *Drosophila* as well as a more targeted mutant analysis in *C. elegans* of the genes within our cluster to examine the phenotypic consequences of their perturbation. In both experimental models, novel candidate genes scored at a similar frequency to known genes within the cluster, without presenting any significant pleiotropic phenotypes. Given the limitations in sensitivity of the assays employed in our screen we conclude that most if not all candidate genes are indeed cilium-linked. We were unable to assess this for the 64 genes not conserved in either worm or fly. However, the plentiful indications of cilium-related functions from large-scale proteomic, expression and localization approaches give us the confidence to extend this conclusion to the entire set of 152 genes within our cluster. The use of *C. elegans* and *Drosophila* as complementary experimental models enabled us to classify novel genes into genes required for cilium biogenesis (the only class conserved in *C. elegans*) or cilium motility, with the former category further subdivided into genes required specifically for compartmentalized cilium biogenesis (scoring in *Drosophila* neurons only) and those required for cilium biogenesis regardless of context (scoring in both *Drosophila* neurons and sperm).

In our downstream characterization we focused on genes required for cilium biogenesis, particularly those involved in the poorly understood early steps surrounding the conversion of centrioles into basal bodies and initiation of transition zone/axoneme assembly. Given the almost total lack of known biogenesis genes conserved in *Plasmodium* where this process must of necessity occur in an IFT-independent manner, we were also interested in any genes required for IFT-independent cilium biogenesis in *Drosophila* sperm. The first set of genes comprises the novel basal body components CABLES1, CCDC6 and TEX9. We found these genes to function in a pathway initiated by the cartwheel component Bld10/CEP135 and also involving the microtubule plus-end tracking protein CEP104. CEP135 is notable for being part of a highly conserved centriole and basal body module first identified by the lab of Monica Bettencourt-Dias, also comprising SASS6 and SAS-4/CENPJ (Carvalho-Santos *et al*., 2010). Yet, unlike SASS6 and SAS-4/CENPJ, CEP135 is not required for centriole assembly *per se*, instead playing a role specifically in the formation of basal bodies (Roque et al., 2012). We see CABLES1, CCDC6 and TEX9 as part of the same centriole to basal body conversion program. While highly conserved, not all components of this program are present in every ciliated species (by our criteria only CABLES1 and CEP104 are universally conserved, including in *Plasmodium*), nor are they necessarily required in every cellular context within the same organism. We suggest that this reflects an underappreciated variability in basal body and ciliary architecture in different species and different tissues within the same species, building on a common core centriole structure (Jana, 2021; Jana *et al*., 2018). A different form of diversification is exemplified by the CFAP97 and CFAP299 family of genes, where multiple paralogs exist in *Drosophila* and vertebrates. It is notable that different family members in many cases have different tissue expression patterns, raising the possibility of both redundancy and functional diversification. Further work will be required to establish the degree to which paralogs can functionally replace each other when expressed in the respective mutant context. When considered as a family, both CFAP97 and CFAP299 function in core (i.e. non-IFT-dependent) cilium biogenesis, with phenotypes in both *Drosophila* sperm and neurons, although only CFAP299 is conserved in *Plasmodium*. As described above, CFAP299 presents a particularly unusual phenotype in sperm, with a failure of membrane deposition around the flagellar axoneme. How this phenotype relates to its function in canonical, compartmentalized cilium biogenesis remains to be established, but it is clear that this is a particularly interesting candidate to follow up on.

Finally, the atypical MAP kinase MAPK15 is of interest as a universally conserved master regulator of ciliogenesis, present in all ciliated species and lost in non-ciliated ones. Such a high degree of conservation stands in contrast to the regulation of centriole assembly, which only in metazoans is driven by the polo-like kinase PLK4/ZYG-1, while core centriolar structural components are universal across eukaryotes (Carvalho-Santos *et al*., 2010). At the same time, the loss of MAPK15 in non-ciliated species argues against pleiotropic functions such as have been proposed for this kinase in vertebrates (Deniz *et al*., 2022). These observations speak to a high degree of evolutionary constraint, potentially reflecting multiple targets within the ciliogenesis pathway. We here identify one downstream effector, the putative ARF/ARL GAP ELMOD, which is likewise highly (but not universally) conserved and which displays a near-identical spectrum of phenotypes to MAPK15 in *C. elegans* and *Drosophila*. The downstream targets for MAPK15 in ciliogenesis beyond ELMOD are currently unclear. While previous work has linked both MAPK15 and ELMOD to ciliary trafficking (Cevik *et al*., 2021; Kazatskaya *et al*., 2017; Turn *et al*., 2022; Turn *et al*., 2021), this clearly cannot explain their role in IFT-independent cilium biogenesis in *Drosophila* sperm and (in the case of MAPK15) *Plasmodium*. It is also notable that no other *C. elegans* gene to date shares their dual phenotype in transition zone assembly and axoneme extension. With both proteins being recruited at the earliest stages of ciliogenesis as centrioles become basal bodies, we suggest that their role is instead in that initial templating step, similar to the role ascribed to TTBK2, a kinase not clearly linked to ciliogenesis beyond metazoans, in the intracellular pathway of ciliogenesis (Goetz *et al*., 2012).

In conclusion, our phylogenetic profiling-based screen has defined what appears to be the core set of genes required for cilium assembly and motility across eukaryotes, a set that includes a substantial number of as yet poorly characterized candidate genes. With many ciliary cluster genes linked to ciliopathies, we believe this dataset to represent an invaluable resource for basic researchers and clinicians alike.

## ACKNOWLEDGEMENTS

We thank members of the Dammermann and Campbell labs for discussions; the *Caenorhabditis* Genetics Center (CGC), the National Bioresource Project for the nematode (NBRP); the Vienna *Drosophila* Resource Center (VDRC), the Bloomington *Drosophila* Stock Center, Clemens Cabernard, Dhanya Cheerambathur, Joe Howard, Jordan Raff and Helen White-Cooper for strains and reagents; Peter Duchek, and Joseph Gokcezade of the IMBA fly house, the Knoblich and Brennecke labs for reagents, fly stocks and help with the *Drosophila* experiments; Daniel Serwas and Nicole Fellner of the VBCF Electron Microscopy facility for help with preparing samples for EM; and Josef Gotzmann and Thomas Peterbauer of the MFPL BioOptics facility for technical assistance. This work was supported by grants P30760 and P34526 from the Austrian Science Fund (FWF) and grant 35376034 from the Austrian Research Promotion Agency (FFG) to A.D., as well as a unidoc PhD fellowship of the University of Vienna to T.Y.S. and a VIPS post-doctoral fellowship of the University of Vienna to J.D..

## AUTHOR CONTRIBUTIONS

J.D. Acquisition of Data (*Drosophila* primary and secondary screen, further analysis), Analysis and Interpretation of Data, Drafting or Revising the Article; T.Y.S. Acquisition of Data (*C. elegans* screen, further analysis), Analysis and Interpretation of Data, Drafting or Revising the Article; B.E. Acquisition of Data (*Drosophila* primary and secondary screen), Analysis and Interpretation of Data; A.S. Acquisition of Data (Bioinformatics); A.D. Conception and Design, Analysis and Interpretation of Data, Drafting or Revising the Article.

## DECLARATION OF INTERESTS

The authors declare no competing interests.

## STAR METHODS

### LEAD CONTACT

Further information and requests for resources and reagents should be directed to and will be fulfilled by the Lead Contact, Alexander Dammermann.

### MATERIALS AVAILABILITY

*C. elegans* and *Drosophila* strains and plasmids generated in this study are available upon request from the Lead Contact.

### DATA AND CODE AVAILABILITY

The published article includes all datasets generated or analyzed during this study.

### EXPERIMENTAL MODEL AND SUBJECT DETAILS

#### *C. elegans* strains and culture conditions

All strains used in this study are listed in Table S3A (primary dyefill screen) and Key Resources Table (further analysis). Mutant strains for the primary screen were obtained from the National BioResource Project (NBRP, *tm* alleles not available at the CGC) or *Caenorhabditis* Genetics Center (CGC, all others). Detailed information on all mutants is available on WormBase (https://www.wormbase.org). Strains expressing endogenous promoter-driven CHE-11:mCherry (Prevo et al., 2015), mCherry:HYLS-1 (Schouteden *et al*., 2015), MKS-6:mCherry (Prevo *et al*., 2015) and OSM-6:GFP (Prevo *et al*., 2015), and the loss of function alleles *ccep-290(tm4927), nphp-4(tm925)* (Schouteden *et al*., 2015), *kap-1(ok676*) and *osm-3(p802*) (Snow et al., 2004) have been previously described. Full gene deletions for *mapk-15*, *ctg-1*, *pgam-5*, and *tag-321* were generated by CRISPR/Cas9-mediated homologous recombination using a plasmid-based protocol (Dickinson et al., 2013). Successful deletion was detected using single worm PCR and confirmed by PCR and sequencing. A full gene deletion for *C56G7.3/elmd-1* was generated by SunyBiotech and verified by PCR and sequencing. A strain expressing mScarlet:PH in neurons was generated by cloning the PH domain of rat PLC1δ1 (Kachur et al., 2008) under the control of the pDyf-7 promoter and unc-54 3’regulatory sequences with an N-terminal mScarlet into the pCFJ151 targeting vector and Mos1-mediated transposition (Frokjaer-Jensen et al., 2008). A strain expressing endogenous promoter-driven GFP:MAPK-15 was similarly generated by cloning the genomic locus plus 5’ and 3’ regulatory sequences with an N-terminal GFP into pCFJ151 and Mos1-mediated transposition. Finally, a strain expressing endogenously GFP-tagged ELMD-1 was generated by SunyBiotech. Dual color strains and strains carrying mutant alleles were constructed by mating. *mapk-15* and *elmd-1* mutants were outcrossed multiple times against wild-type prior to analysis/introduction of fluorescent markers. All strains were maintained at 20-23°C.

#### Drosophila melanogaster stocks and husbandry

All strains used in this study are listed in Table S3A (primary and secondary screens) and Key Resources Table (further analysis). Strains were obtained from Helen White-Cooper (Bam-GAL4 driver line), the Bloomington *Drosophila* Stock Center (BDSC, w[1118] allele, Tub-GAL4, Elav-GAL4 and Nanos-GAL4 driver lines, as well as TRiP RNAi fly stocks) and Vienna *Drosophila* Resource Center (VDRC, GD and KK RNAi fly stocks). Detailed information for these strains is available on FlyBase (http://flybase.org), as well as on the websites of the two stock centers (BDSC, https://bdsc.indiana.edu, VDRC, https://stockcenter.vdrc.at/control/main). Cep135 mutants were obtained from the Cabernard lab. Cep104 and Cables1 mutants were generated by Flp-FRT recombination between two transposon elements flanking its coding sequence (Parks et al., 2004). In the case of Cep104 (CG10137) these were PBac{RB}CG10137e00472 and PBac{WH}CG10137f04546, in the case of Cables1 (CG6191) PBac{RB}CG6191e03779 and P{XP}CG6191d03529. To obtain complete deletions, males carrying one element were mated with females carrying a FLP recombinase transgene. Progeny males carrying both the element and FLP recombinase were then mated to females carrying the other element. After 2-3 days, parents were transferred to a new vial and progeny subjected to a 1h heat shock by placing the vials into a 37°C water bath. We subjected the vials to daily 1h heat shocks for another 4 days. Progeny were then raised to adulthood, males collected and crossed to females containing marked balancer chromosomes. Single progeny males were then crossed individually to virgin females to generate additional offspring for PCR confirmation analysis and to balance the stocks in an isogenic background. A full deletion of CFAP299 (CG8138) was generated by CRISPR/Cas9-mediated genome editing (Port et al., 2014). Two guideRNAs were cloned in the pCFD4 vector, one for the start and one for the end of the coding sequence. Co-injection with a DsRed rescue plasmid (pHD-DsRed-attP plasmid (Gratz et al., 2015)) with 1000bp flanking sequence on either side of the gene were carried out in vas-Cas9 flies (BDSC #55821) by BestGene Inc.. DsRed-containing flies were screened by PCR to ensure proper targeting. The GFP-fusion to Cep135/Bld10 under the control of the Ubiquitin promoter (Roque *et al*., 2012) was a gift of the Raff lab. Tex9 (CG4681) GFP was a FlyFos construct (Sarov et al., 2016) from the VDRC (#318643). All other GFP strains were generated as described previously (Stevens et al., 2010) using Ubq-GFP plasmids for N and C-terminal tagging. In brief, P element–mediated transformation vectors containing GFP-fusions to Cables1 (CG6191), CCDC6 (CG6664), Cep104 (CG10137), CFAP97D1 (CG14551), Elmod (CG10068), and Mapk15 (CG32703) were generated as follows: The complete coding region of each protein was amplified from cDNA with att sites at either end for Gateway cloning (Invitrogen). These fragments were inserted into the Gateway pDONR Zeo vector. The pDONR vectors were then recombined with Ubq plasmids (Peel et al., 2007), with each coding sequence placed in-frame with GFP at the N or C terminus. The transgenic lines were generated by BestGene Inc.. The Ubq promoter drives moderate expression in all tissues (Lee et al., 1988). Expression of N and C-terminal constructs was examined in testes and neurons.

### METHOD DETAILS

#### Phylogenetic profiling

Genomes were selected as widely as possible from all branches of the eukaryotic tree, taking care to avoid overrepresenting certain branches (See Table S1A). In addition, a number of bacterial and archaeal genomes were included as outgroups. Only completely sequenced and well annotated genomes were chosen by confirming the presence of universally conserved genes. Starting point for our analysis was the human proteome, obtained from UNIPROT on 7 Feb 2014. Datasets were generated by performing a reciprocal BLASTp analysis for each human protein coding gene against the proteomes of each of the 159 genomes in our analysis. The BLAST-algorithm used was NCBI BLAST 2.2.28, build 12 Mar 2013. If a hit was found above the EXPECT threshold of 0.1, the highest scoring protein sequence in this proteome was then run against the human proteome. If this reciprocal BLAST returned a protein of the same GeneID, a reciprocal hit was established for this protein, in that species, represented by a 1 in the output dataset. Otherwise a 0 was reported for this protein-species test. The resulting binary matrix was then used as input for hierarchical clustering in Gene Cluster, using the Euclidean distance metric and average linkage clustering for both genes and species on the entire dataset comprising 21732 human protein coding genes and 159 species. Within the clustering output viewed in Java TreeView a group of genes was identified with a similar phylogenetic inheritance pattern centered around the core centriolar component CENPJ/SAS-4 (Kirkham et al., 2003). This cluster appeared quite distinct, such that loosening similarity constraints would only gradually grow the cluster until a final size of 386 genes was reached. As detailed elsewhere in this manuscript, this cluster was highly enriched in known ciliary genes and hence was chosen for further analysis. To evaluate the robustness of our method, we repeated the hierarchical clustering with reduced numbers of species (25%, 50% or 75% fewer), with genomes removed equally from all branches of the eukaryotic tree (see Table S1A). The resultant clusters of genes centered around CENPJ were then compared with the original, full ciliary cluster and any differences in gene number and identity noted.

#### Identification of *C. elegans* and *Drosophila* orthologs

*C. elegans* and *Drosophila* orthologs of genes within the ciliary cluster were manually identified by reciprocal BLAST analysis using the human protein as the starting point. Where direct comparisons failed to identify a clear homolog, indirect searches were performed using less divergent related species as intermediates. To resolve gene duplications in one or other species, multiple sequence alignments were generated using MUSCLE within Jalview (http://www.jalview.org), with average distance phylogenetic trees constructed also in Jalview using the BLOSUM62 matrix.

#### Literature searches and database analyses

For each of the 386 genes in the cluster we conducted a comprehensive literature and database analysis, including any information on their putative orthologs in non-vertebrate experimental models. For literature searches in PubMed we employed all alternative names and classified each gene based on its level of functional characterization overall and any previously reported link to cilia. We further set out to provide a short description of its proposed function, with broader and more specific classifiers (e.g. Centriole/cilium biogenesis, IFT-A complex), including references to the relevant literature with PubMed identifiers. Any potential disease association relevant to ciliary function was reported separately, compiling reports from PubMed and OMIM (https://omim.org). Database analyses were conducted by retrieving the relevant datasets from the original source publication or website as indicated. To ensure proper comparisons, all identifiers were converted into Ensembl Gene IDs and their presence in the full human proteome dataset verified manually. The same was done with the sets of genes associated with specific centrosomal/ciliary functional modules, compiled from recent reviews and primary research publications. Classification of genes into predicted cilium biogenesis/motility genes was based on degree of conservation in ciliated/non-ciliated species, presence in species with non-motile cilia, species with a reduced complement of ciliary genes and *Plasmodium*, with cut-offs chosen manually based on the observed pattern for known ciliary genes (see Table S2D).

#### *C. elegans* primary screen

Dye-fill assays were performed on L4 stage animals. For each condition, ∼100 worms were picked into 250µl of M9 0.1% Triton, washed 3x with M9 0.1% Triton, 1x with M9, and then incubated at room temperature under dark conditions for 1h in 0.2% DiI (Invitrogen) in M9. After incubation, worms were destained for at least 30min on a seeded NGM plate before analysis. Dye accumulation in cell bodies of amphid and phasmid neurons was scored on a Zeiss Axio Imager Z2 microscope equipped with a 63x 1.4NA Plan Apochromat objective and Lumencor SOLA SE II light source. For illustration purposes, 0.2µm z-series were acquired DeltaVision 2 Ultra microscope, equipped with 7-Color SSI module and sCMOS camera and controlled by Acquire Ultra acquisition software (GE Healthcare). Image stacks were deconvolved using Softworx and then imported into Fiji for post-acquisition processing.

Dyefill phenotypes were assessed using the following scoring system: Amphids: 0 = 11.5-12 dye-filling neurons, 1 = 11-11.49, 2 = 10-10.9, 3 = 6-9.9, 4 = 0-5.9, Phasmids: 0 = 3.85-4 dye-filling phasmids, 1 = 3.75-3.84, 2 = 3.5-3.74, 3 = 0.25-3.49, 4 = 0-0.24. The strongest phenotype of amphids and phasmids was reported as the overall phenotype.

#### Further analysis of amphid and phasmid neurons in *C. elegans*

Live imaging of embryos and larvae was performed using a 60x 1.42NA Plan Apochromat objective on a DeltaVision 2 Ultra microscope, equipped with 7-Color SSI module and sCMOS camera and controlled by Acquire Ultra acquisition software (GE Healthcare). Worms were anesthetized using 10mM tetramisole in M9 for 5min before being mounted on 2% agarose pads and imaged at room temperature. Embryos were mounted directly on agarose pads. 0.2µm GFP/mCherry z-series as well as single plane DIC images were collected for the amphid and phasmid neurons closest to the objective. For each condition and worm stage, at least 10 worms/embryos were examined. Image stacks were deconvolved using Softworx and then imported into Fiji for post-acquisition processing. For strains expressing OSM-6:GFP, 0.5µm z-series were acquired on a Zeiss Axio Imager Z2 microscope (described above) using a Photometrics CoolSNAP-HQ2 cooled CCD camera controlled by ZEN 2 blue software (Zeiss).

Quantitation of phasmid dendrite lengths (from cell body to ciliary base) and ciliary length (from base to tip) were performed in Fiji using the Segmented Line tool. For each condition, a minimum of 20 worms were examined.

#### Transmission electron microscopy of *C. elegans*

L4-stage worms were prepared using chemical fixation following the protocol described in (Serwas and Dammermann, 2015). Worms were fixed in 2.5% glutaraldehyde in cytoskeleton buffer (CB, 100mM methyl ester sulfonate, 150mM NaCl, 5mM EGTA, 5mM MgCl_2_, 5mM glucose in ddH_2_O, pH 6.1) overnight at 4°C. Samples were then washed 3x in 1xCB and post-fixed for 30min in 0.5% osmium tetroxide in CB. Fixed worms were then washed 3x in CB and 1x in ddH_2_O. Finally, samples were dehydrated for 15min each in 40%, 60%, 80%, 2x in 95% and 3x in 100% acetone. Samples were embedded in Agar100 resin after fixation and dehydration. 70nm serial sections were prepared onto Cu 100mesh grids with Formvar film, then post-stained with aqueous uranyl acetate and lead citrate and examined with a Morgagni 268D microscope (FEI) equipped with an 11-megapixel Morada CCD camera (Olympus-SIS) and operated at 80 kV.

#### *Drosophila* primary screen

To deplete specific target genes in the whole fly or specific tissues three different RNAi crosses were set up. We used Tub-GAL4 as a ubiquitous driver. Tissue-specific drivers used were: a combination of Bam-GAL4 (Chen and McKearin, 2003) and Nanos-GAL4 (gift of Helen White-Cooper) for germline depletion and Elav-GAL4 (Lin and Goodman, 1994) for neuronal depletion. In each case we crossed 5-8 virgins to 4 males from the respective RNAi strain (KK and GD: (Dietzl et al., 2007), TRiP: (Ni et al., 2008)) in standard cornmeal-agar vials. Vials were kept for 3 days at 25°C, 60% relative humidity, 12/12 light-cycle. On the end of the third day we discarded the parents and vials were moved to 29°C, 60% relative humidity, 12/12 light cycle.

To test male fertility we collected 5 males. Males were crossed to 5 w-virgin females (3-7 days old) in a new vial. As females do not lay eggs in the direct neighborhood of yeast granules, we used standard cornmeal vials without dry yeast to eliminate bias due to differences in yeast-free surface availability. Fertility test crosses were kept at 29°C to strengthen knock-down efficiency. After 3 days adults were removed. Uncoordination, wing posture and flight was scored in Elav and Tubulin crosses. Male fertility was scored in Bam/Nanos and Tubulin crosses. Viability was scored in all crosses. The strongest phenotype was reported from the different RNAi lines tested.

Viability was scored comparing the number of dead larvae, dead pupae and dead pigmented wing pharate adults to the overall progeny number. Viability was scored for all three stages. Few or no progeny was scored as larval lethality. To confirm early lethal phenotypes crosses were repeated two to three times. The following scores were used: 4 = complete lethality, 3 = strong lethality with a few survivors, 2 = half of the progeny died at a given stage, 1 = dead individuals, but less than half of all offspring, 0 = no or few (<3) dead individuals were observed. For sperm and neuron-specific RNAi scores reported are a combination of pupal and pharate lethality.

To test male fertility we counted pupae 13 days after setting up the test cross with w-when all progeny already pupariated. Control crosses under these conditions produced on average 120 pupae. Pupa numbers were scored using the following scheme: 0 = 90 pupae or more, 1 = between 89 and 60, 2 = between 59 and 30, 3 = between 29 and 10, and 4 = less than 10.

Adult flies were analysed using the following classification system to score uncoordination and flight. Uncoordination was scored as follows: 4 = Strong uncoordination with all adults sticking to the surface of the food, 3 = Uncoordinated flies at the bottom of the vial unable to climb up the vial. Weak uncoordination (2 and 1) was scored using a flight test. 20-30 flies were put into a fresh vial and after 1h of recovery we tested the ability to fly away from the rim of the vial. Flight was scored as follows: 2 = more than half of the flies were unable to fly, 1 = less than half of the flies were unable to fly (> 3 no fliers). As wing defects affected flying ability, only flies without wing defects were used for flight test.

Wing posture defects scored were lowered wing blades, extended wings and completely erected or completely dropped wings. We scored the frequency of these phenotypes (> 20 counted): 4 = all individuals affected, 3 = more than half affected, 2 = half of flies affected, 1 = several individuals affected, 0 = No or few individuals affected (<3).

#### *Drosophila* secondary screen - Analysis of male germline defects

Fly strains were crossed as in the primary screen. 0-1 day-old males were dissected in PBS. The tip of the abdomen with testis attached was placed in 4% formaldehyde in PBS for 25min and washed 3×5min in 0.1% PBS-Triton-X. Testes were transferred into a droplet of staining solution of 5µg/ml Hoechst 33342 and a 1:50 dilution of Alexa 568 phalloidin in PBS for two hours. Samples were washed overnight at 4°C in 1ml of PBS. Three pairs of testes were mounted in mounting medium (2% w/v n-propyl gallate in PBS/ 90% Glycerol) on multi-well slides and covered with a coverslip. Slides with at least 3 pairs of testes per genotype were imaged on a Zeiss LSM 700 confocal laser scanning microscope equipped with 25x Plan-Apochromat oil immersion objective. Image stacks of approximately 20 z-planes were acquired at 4µm increments for each of the 6 testes. For a complete testis image stack 4 z-stacks were stitched together using ZEN software (Zeiss). Motility of sperm was tested by dissecting testes with seminal vesicle attached in a drop of Schneider media. The seminal vesicle was pierced with a sharp tungsten needle and sperm motility imaged immediately using dark field settings on a Zeiss Axio Imager Z2 microscope equipped with a 10x Plan-Neofluar objective at 50 frames/s.

To classify defects we distinguished six stages of spermiogenesis (Tates, 1971): Gonialblast stage (Tates stage 1,2,3), spermatocyte stage (large masses of heterochromatin at the periphery of the nucleus, Tates stage 4,5,6), spermatid stage with round nucleus (Tates stage 10-14), spermatid stage with elongated nucleus (Tates stage 17-18), stacked/pinhead sperm/spermatid stage (Tates stage 19,20). Additionally, we scored the number and shape of the actin cones during sperm individualization (marked by phalloidin-positive membrane investment cones). Defects were scored for each stage using the following scale: 0 = normal morphology, 1 = weak deviation (occasionally observed in negative controls), 2 = stronger effect but normal morphology also observed, 3 = all or vast majority of nuclei/investment cones for a given stage show aberration, 4 = missing stage. Motility was assessed visually based on the number of motile sperm and frequency of flagellar beat compared to wild-type sperm. Absence of sperm from the seminal vesicle was also noted.

#### *Drosophila* secondary screen - Analysis of defects in chordotonal neurons by DIC microscopy

Chordotonal organ samples were taken from P9 stage male puparia (Bainbridge and Bownes, 1981). Puparia were removed from the side of the vial with a pair of forceps and glued ventral side down to a slide covered with double-sided adhesive tape. The operculum was removed and puparial case cut open along the dorsal mid-line. Pharate adults were taken out of the case and glued dorsal side down on the tape. Front legs were torn off and mounted on a slide in a droplet of PBS and covered with a cover slip. Gentle pressure was applied on the coverslip to expel the chordotonal organ from the femur. At least three chordotonal organs from three different individuals were imaged for each genotype on a Zeiss Axio Imager Z2 microscope equipped with 63X Plan-Apochromat oil immersion objective in DIC modus. For each organ a stack of 20 z planes were acquired at 0.5µm increments.

The primary unit for quantification was the scolopidium, which contains two ciliated neurons. Scolopidia are organized in three groups (1, 2 and 3, (Shanbhag et al., 1992)). We only used scolopidia from groups 2 and 3. For morphological, numerical quantification and dilation distance measurements three chordotonal organs were used from three different individuals. Screening for dilation defects was done on all images for a given genotype. Defects in scolopidium morphology were observed and scored in the following way: ‘No cilium’ – cilium not detected, ‘Short cilium’ – tip does not reach the cap cell, ‘Regressive cilium’ – thinner, bent ciliary shaft and ‘Vesicles in scolopidial space’. To measure the strength of each phenotype the ratio of morphologically defective scolopidia compared to the total number of scolopidia of a given chordotonal organ was calculated. A gene was scored as abnormal if more than 20% of scolopidia contained neurons that were morphologically defective. To analyse the presence or absence of cilia, we counted the number of scolopidia with no cilium, 1 cilium, 2 cilia or more than 2 cilia. In addition, we assessed morphological defects and mispositioning of the dilations.

#### *Drosophila* secondary screen - Analysis of defects in chordotonal neurons using Sas-4 and NompC staining

Leg chordotonal organs were dissected from 36h old male pupae pupae. Fixation and staining were done as previously described for testes (Martinez-Campos et al., 2004). Briefly, legs were dissected in PBS. Leg chordotonal organs were placed between slide and coverslip and pushed out of the cuticle by applying gentle pressure. Slides were then snap-frozen in liquid nitrogen. After recovering slides and removing the coverslip, samples were incubated in ice-cold methanol for 5min, followed by ice-cold acetone for 2min. Samples were then washed twice with PBS 0.5% Triton X-100, followed by blocking in PBS 0.1% Triton X-100 1% BSA for 30min. Primary antibodies for NompC and Sas-4 were diluted 1:500 in blocking solution and added to the samples and incubated overnight at 4C. The next day samples were washed twice with PBS before addition of secondary antibody in PBS for 2h at room temperature. After washing once with PBS, Hoechst 33342 was added for 10min. Samples were air-dried and mounted with mounting medium. Defects for Sas-4 staining (absence) and NompC staining (absence, misposition, redistribution) were scored. 3 different legs were analysed and the average defect per scolopidium reported.

#### Detailed analysis by immunofluorescence microscopy in *Drosophila*

For detailed examination of sperm centrioles and cilia, testes were dissected from 72h-old male pupae. Fixation and staining were done essentially as described above for chordotonal organs. After dissection, testes were transferred onto a microscope slide and cut with a tungsten needle before placing on a coverslip. Immunofluorescence samples were analyzed on a Zeiss LSM980 scanning confocal microscope equipped with an Airyscan unit. Stacks of 0.75μm slices were acquired with a 63x 1.4NA Plan Apochromat lens using single channel mode to avoid cross-illumination. Airyscan images were acquired for selected centrioles using 0.25μm slices and 10x electronic magnification. Maximum intensity projections of Z stacks prepared in ImageJ were used for image analysis and panel preparation.

#### Transmission electron microscopy of *Drosophila*

To examine ultrastructure of chordotonal organs, legs from 36h old pupae were cut off with microscissors and fixed using a mixture of 2% glutaraldehyde and 2% paraformaldehyde in 0.1mol/l sodium phosphate buffer, pH7.2 for 2h in a desiccator at room temperature and then overnight on a rotator at 4°C. Legs were then rinsed with sodium phosphate buffer, post-fixed in 2% osmium tetroxide in buffer on ice, dehydrated in a graded series of acetone on ice and embedded in Agar 100 resin. 70nm sections were cut and post-stained with 2% uranyl acetate and Reynolds lead citrate. Sections were examined with a Morgagni 268D microscope (FEI, Eindhoven, the Netherlands) operated at 80kV. Images were acquired using an 11 megapixel Morada CCD camera (Olympus-SIS).

To examine sperm flagellar ultrastructure, late pupal testes from third instar larvae were dissected in PBS and fixed using 2.5% glutaraldehyde in 0.1mol/l sodium phosphate buffer, pH7.2 for 1h at room temperature. Samples were then rinsed with sodium phosphate buffer, post-fixed in 2% osmium tetroxide in dH2O on ice, dehydrated in a graded series of acetone and embedded in Agar 100 resin. 70nm sections were then cut and processed as above.

### STATISTICAL ANALYSIS

All error bars are mean with standard deviation. Unpaired t-tests were conducted using GraphPad Prism to compare samples in a specific experiment. *, **, *** represent P-Values of <0.05, 0.01 and 0.001, respectively. Unless otherwise stated, tests are comparing indicated condition to control.

## KEY RESOURCES TABLE

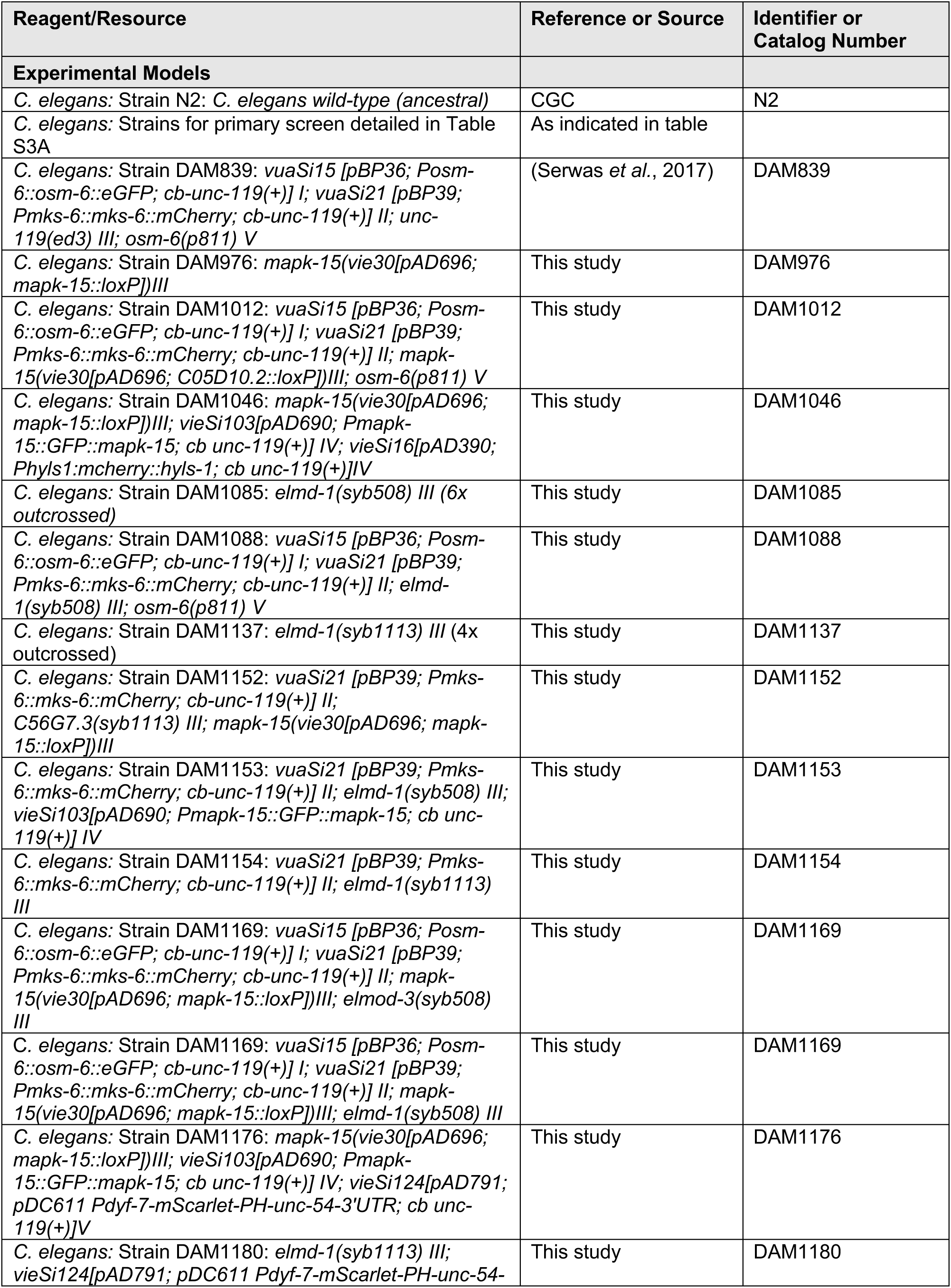
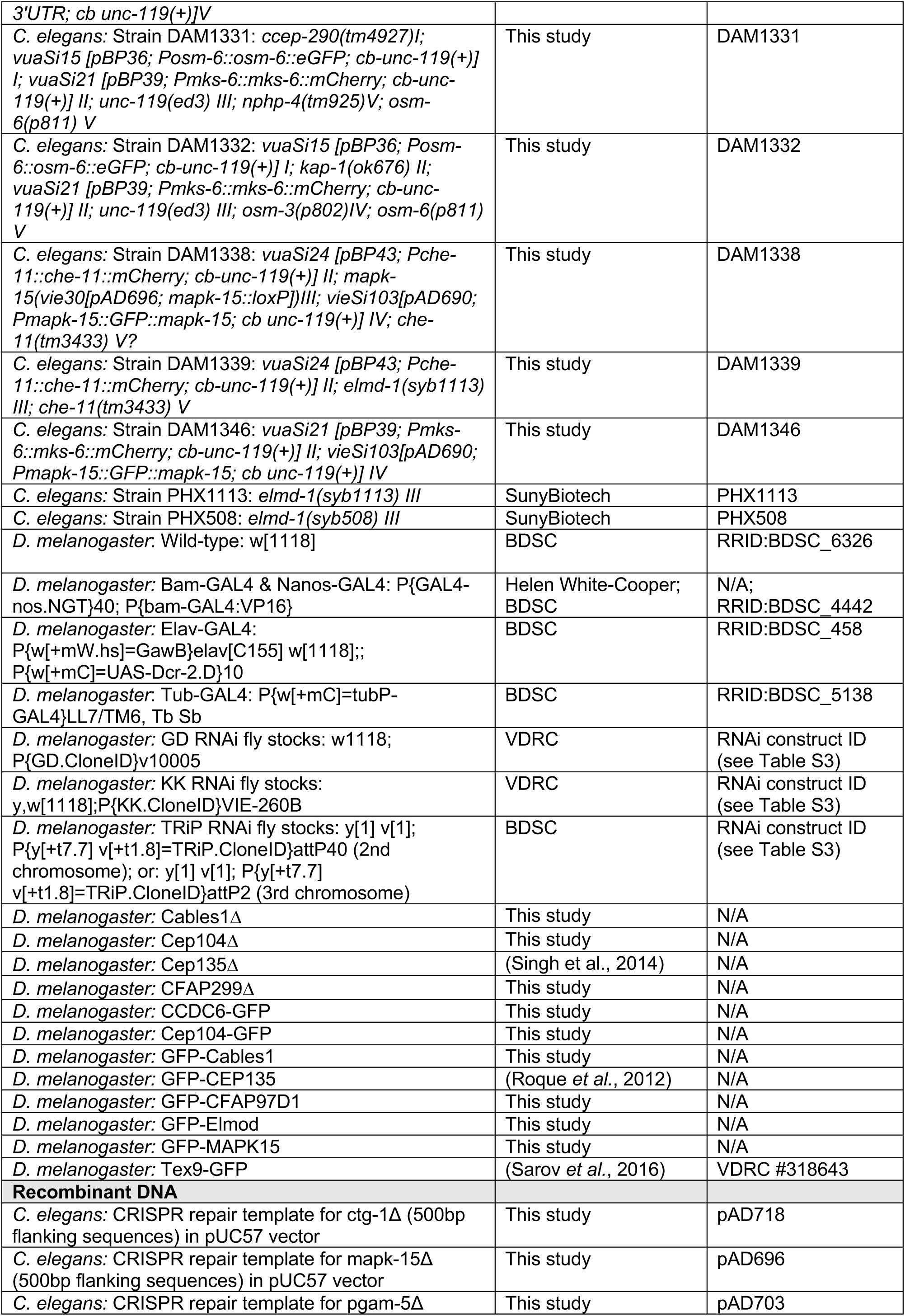
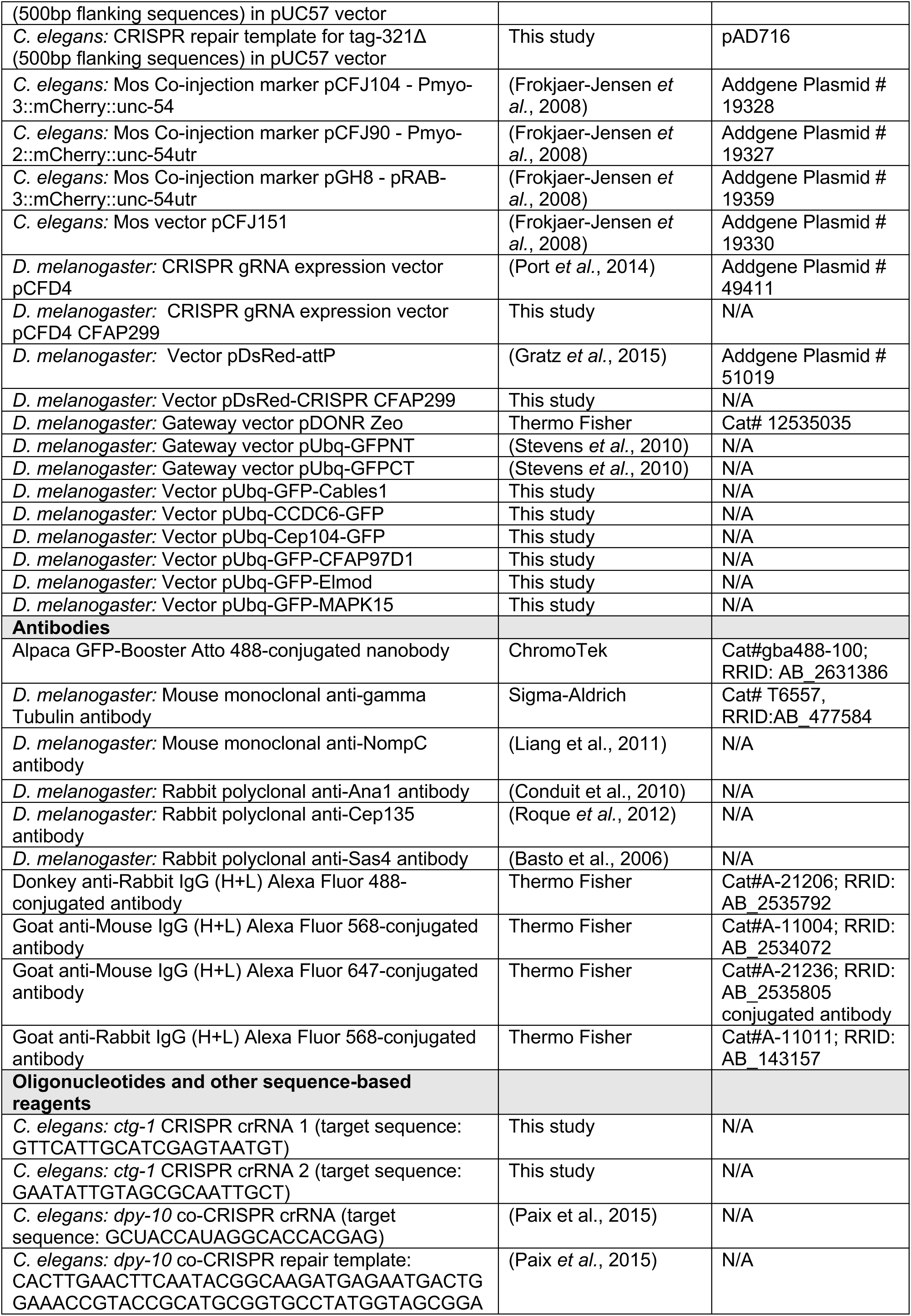
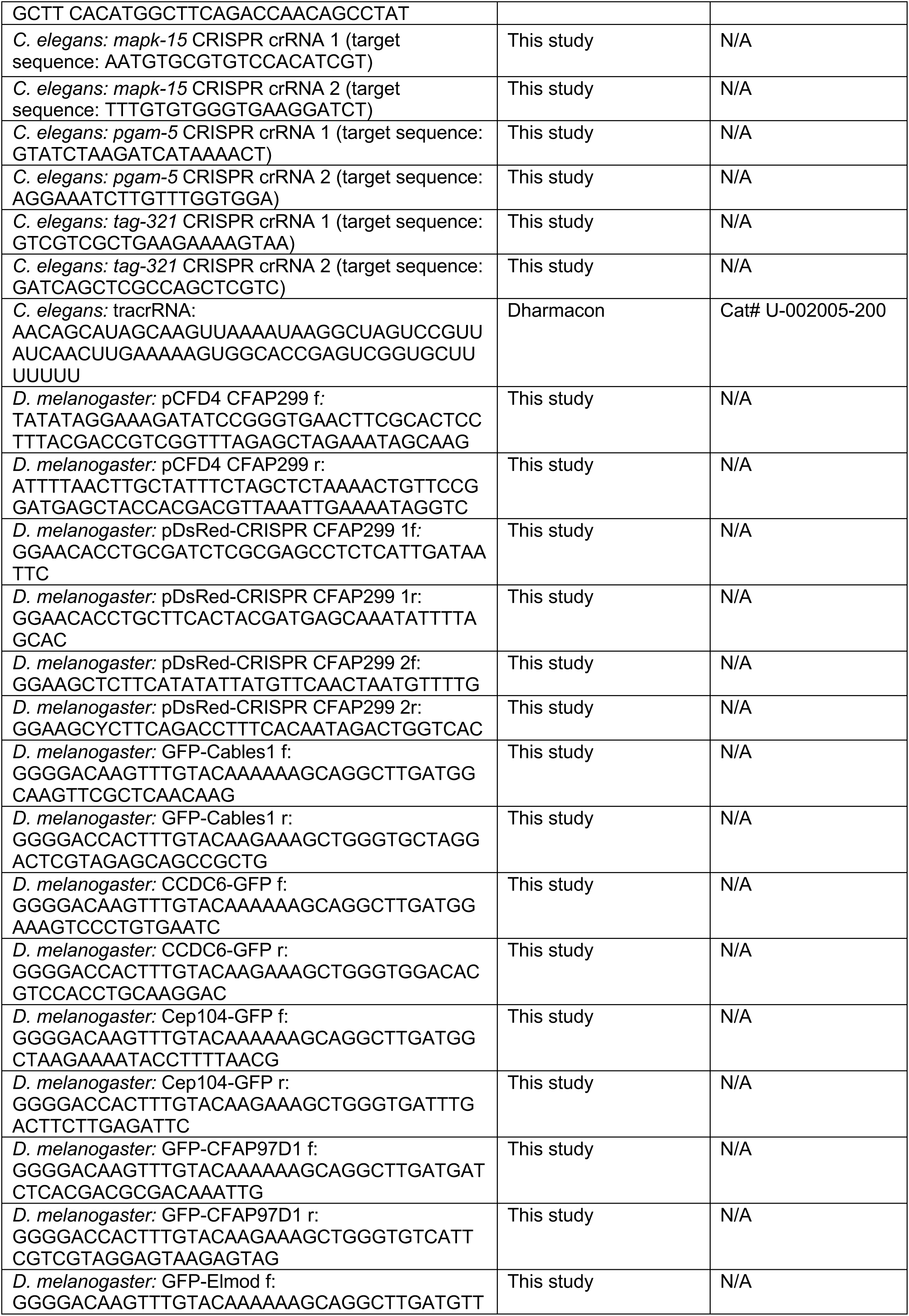
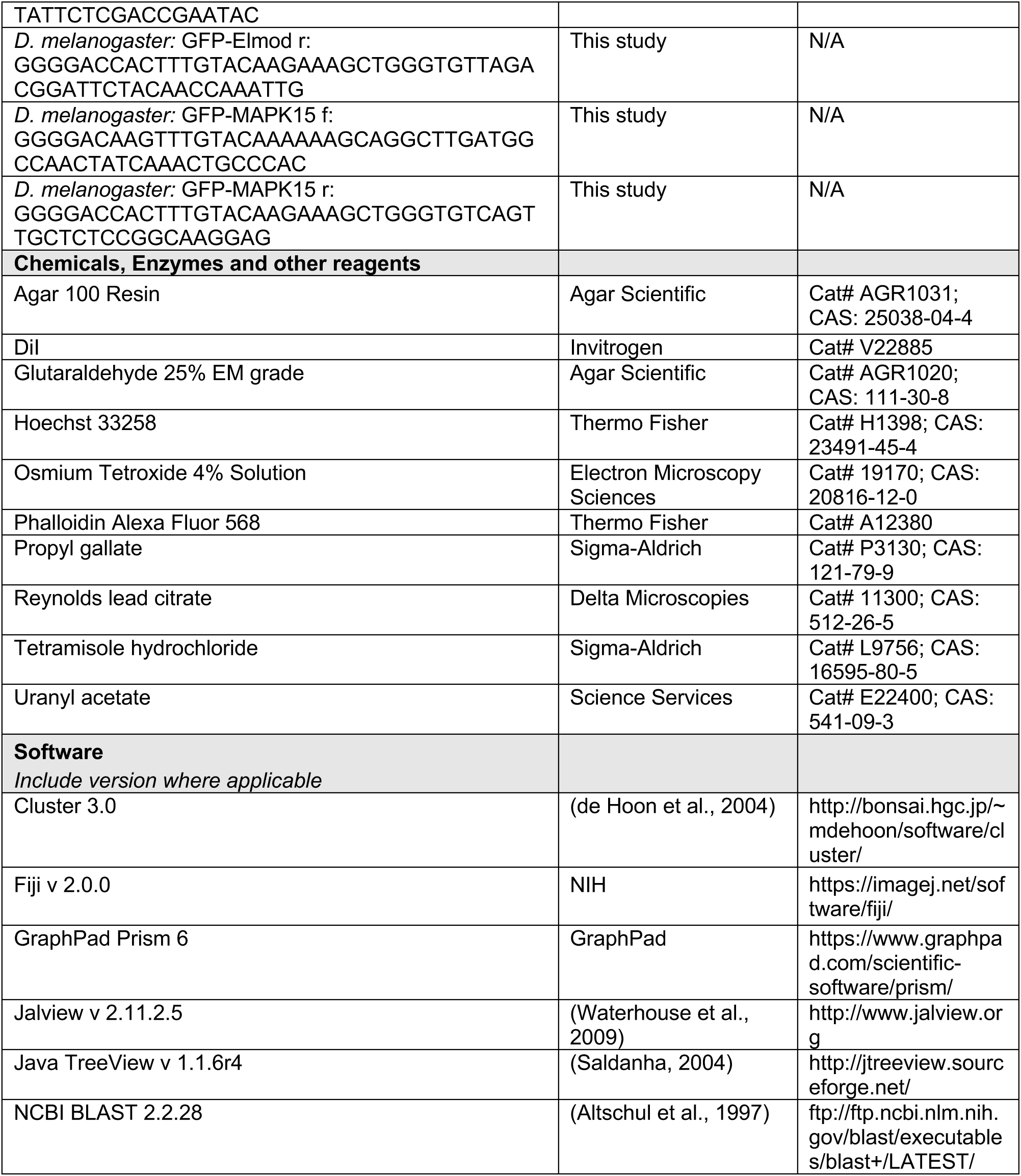

## SUPPLEMENTAL FIGURE LEGENDS

**Figure S1:**
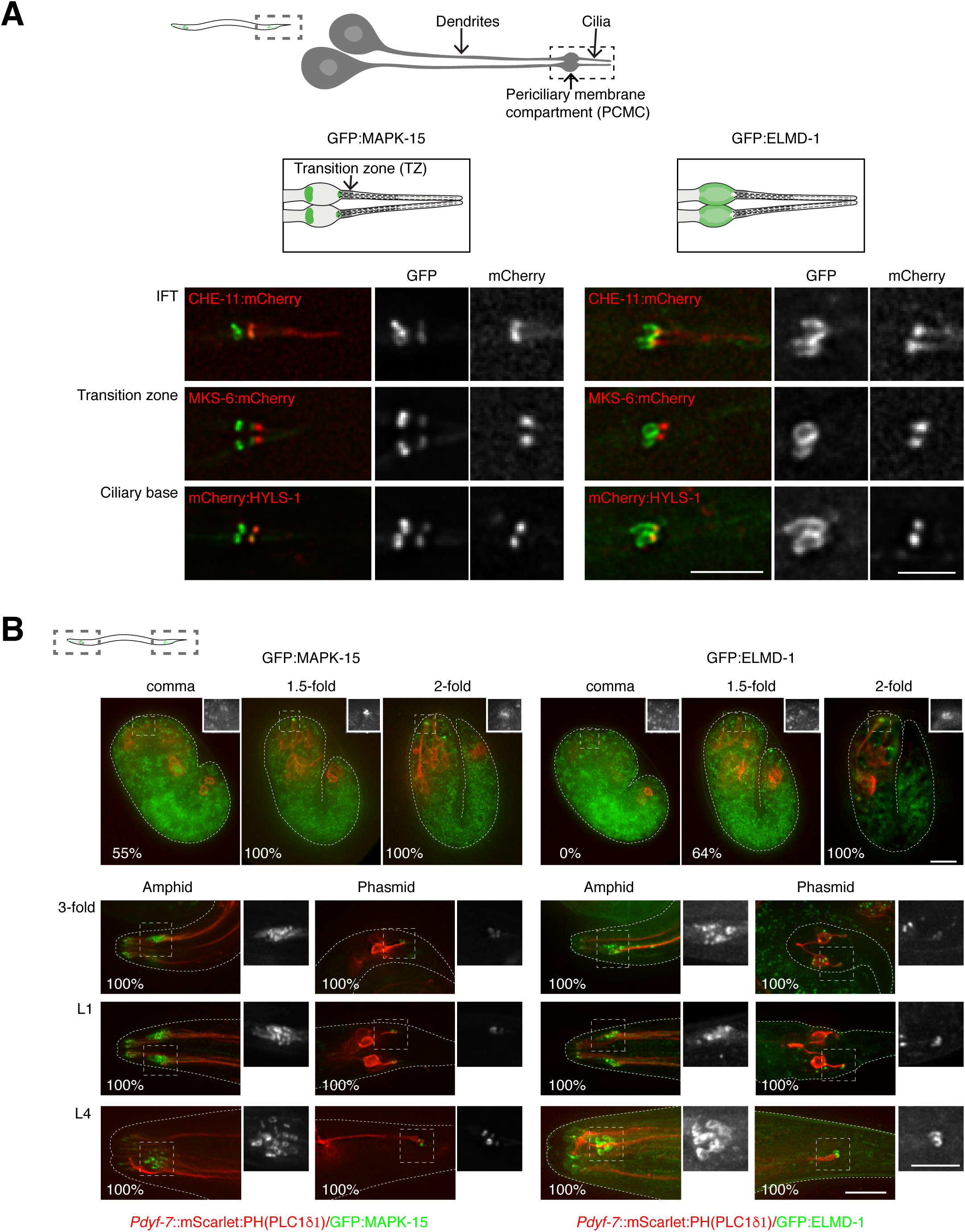
Further characterization of *C. elegans* MAPK-15 and ELMD-1. (**A**) Detailed examination of MAPK-15/ELMD-1 localization in *C. elegans* phasmid neurons. Endogenous promoter GFP:MAPK-15 co-localizes with the basal body marker HYLS-1 and accumulations of the IFT marker CHE-11, just proximal to the transition zone marked by MKS-6. A second population of MAPK-15 localizes to the adhesion belt at the proximal end of the periciliary membrane compartment. Endogenously GFP-tagged ELMD-1 localizes to the entire periciliary membrane compartment. (**B**) Analysis of MAPK-15/ELMD-1 recruitment during *C. elegans* embryogenesis. MAPK-15 signal first becomes detectable in post-mitotic sensory neurons marked by expression of a plasma membrane marker at the comma stage of embryogenesis (430 min after fertilization, (Sulston et al., 1983)), with ELMD-1 signal following shortly thereafter at the 1.5-fold stage (460 min), both proteins localizing to the distal tip of the elongating dendrite where the cilium will eventually form (Nechipurenko et al., 2017; Serwas *et al*., 2017). Neither GFP fusion is detectable at earlier stages of development or in other tissues of the worm. The hazy fluorescence signal in late stage embryos is due to autofluorescence. Scalebars are 10µm (A, B), 5µm (A, B, insets).

**Figure S2:**
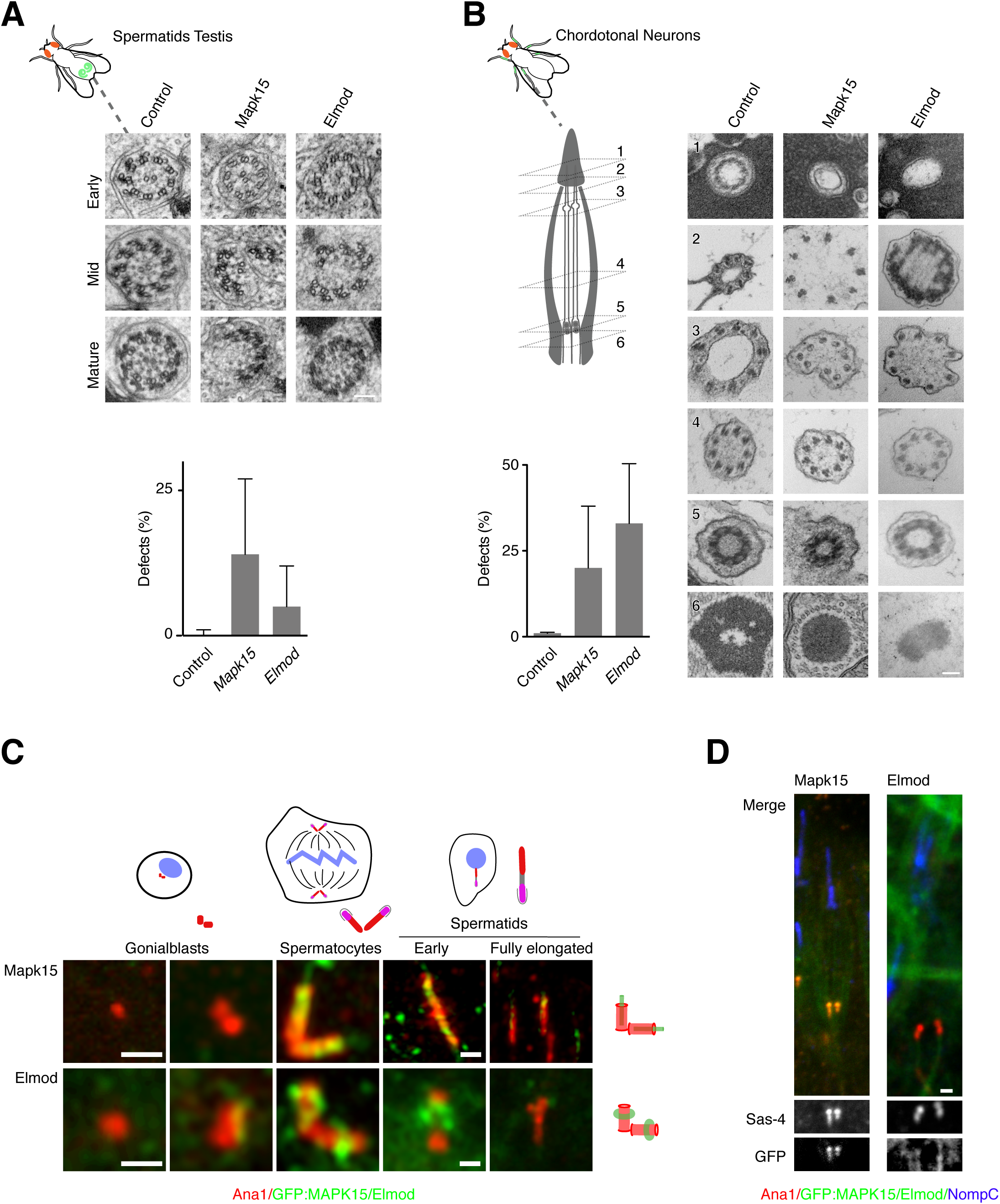
Characterization of Mapk15 and Elmod in *Drosophila*. (**A**, **B**). RNAi-mediated depletion of MAPK15 and ELMOD in *Drosophila* sperm (A) and chordotonal neurons (B) results in defects in ciliary ultrastructure in both cellular contexts, with shortened and broken axonemes in sperm and neurons, respectively. At least 3 animals examined per condition. (**C**) GFP fusions to MAPK15 and ELMOD do not localize to centrioles marked by Ana1, but are recruited to maturing basal bodies during *Drosophila* spermatogenesis. (**D**) MAPK15 and ELMOD co-localize with the centriolar marker Sas-4 on mature basal bodies in chordotonal neurons. Scalebars are *lOOJLm* (A, B), lJLm (C, D), Error bars in (A, B) are standard deviation.

## SUPPLEMENTAL TABLES

**Table S1: Phylogenetic profiling.** (**A**) Genomes used in phylogenetic profiling analysis, including source and taxonomy classification according to (Schoch et al., 2020) and (Baldauf, 2008). Ciliary status assessed *a posteriori* based on presence of known cilium biogenesis/motility genes. Also marked in the table are the genomes included when testing the robustness of ciliary cluster identification by reducing the genome set by 25%, 50% and 75%. (**B**) Hierarchical clustering output including original UniProtKB identifier, gene name and description, as well as length of sequence chosen in starting human proteome dataset. Presence and absence pattern across 159 genomes based on reciprocal best match criterion indicated by 1s and 0s. Marked within the ciliary cluster (1-386, found within the clustering output at 3470-3855) are those genes that continue to be identified when the starting set of genomes for hierarchical clustering was reduced as described in (A).

**Table S2: *In silico* analysis of ciliary cluster.** (**A**) Datasheet summarizing results of literature analysis for each ciliary cluster gene, including any information for putative orthologs in commonly used non-vertebrate experimental models. For PubMed analysis, a score was given based on the total level of characterization (regardless of whether the gene is considered cilium-linked or not) and level of characterization of any potential ciliary function (0 no information, 1 some indications of function, 2 confirmed function, 3 clearly established molecular mechanism). Any gene scoring 2 or 3 for ciliary function was considered ‘known’, with a detailed description including classification into broader functional categories and relevant PubMed IDs provided in subsequent columns. Any gene scoring 0 or 1 for ciliary function was considered ‘unknown/novel’, regardless of the level of characterization for other, unrelated cellular functions. Disease association is based on OMIM (https://omim.org) as well as PubMed analysis, with relevant identifiers provided. Only (potential) cilium-related disorders were considered in this table. SYSCILIA gold standard (SCGSv2) classification was retrieved from the associated recent publication (Vasquez *et al*., 2021), as were CiliCarta scores (van Dam et al., 2019). Cildb V3.0 number of ciliary evidences (Arnaiz et al., 2014) were collected for each human gene from http://cildb.i2bc.paris-saclay.fr. (**B**) Lists of genes associated with specific centrosomal/ciliary functional modules and their presence or otherwise within the ciliary cluster, compiled from recent reviews and primary research publications: Centriolar and PCM components (Jana, 2021); distal and sub-distal appendages (Tischer et al., 2021); transition zone (Garcia-Gonzalo and Reiter, 2017), IFT kinesin and IFT-A/B (Klena and Pigino, 2022), IFT dynein, inner and outer dynein arms, DNAAFs and MIPs (Braschi et al., 2022), the BBSome (Wingfield et al., 2018), ciliary membrane trafficking (Zhao et al., 2023), the N-DRC (Bower et al., 2013), radial spokes 1 and 2 (Gui et al., 2021), radial spoke 3 (Bazan et al., 2021; Urbanska et al., 2015), central apparatus (Gui et al., 2022), and centriolar satellites (Prosser and Pelletier, 2020). (**C**) Comparison of ciliary cluster with the original comparative analyses performed in 2004 (Avidor-Reiss *et al*., 2004; Li *et al*., 2004) and the output of the more recent phylogenetic profiling studies (Dey *et al*., 2015; Li *et al*., 2014; Nevers *et al*., 2017). Gene names for (Avidor-Reiss *et al*., 2004) obtained from original publication, with human orthologs obtained based on reciprocal BLASTp best match criterion, and (Nevers *et al*., 2017) for all others. (**D**) Comparison of predicted function of ciliary cluster genes based on phylogenetic pattern of conservation and actual function (where known). Predicted function based on degree of conservation in ciliated/non-ciliated species, presence in species with non-motile cilia, species with a reduced complement of ciliary genes and Plasmodium (an outlier with a highly reduced complement of ciliary genes), as indicated in the table. Actual reported function derived from (A). (**E**) Presence of ciliary cluster genes in large-scale proteomic, expression and localization screens performed in different experimental models, obtained from the following sources: (1) Proteomics: *Chlamydomonas* ciliary proteome (Pazour *et al*., 2005), from http://chlamyfp.org/ChlamyFPv2/index.php; trypanosome ciliary proteome (Broadhead et al., 2006); *Paramecium* ciliary proteome (Arnaiz et al., 2009), from http://cildb.i2bc.paris-saclay.fr; sea anemone, choanoflagellate and sea urchin ciliary proteomes (Sigg et al., 2017); (2) Expression: *Chlamydomonas* upregulated on deflagellation (Albee et al., 2013); zebrafish Foxj1-induced genes (Choksi et al., 2014); *Xenopus* Rfx2 target genes (Chung et al., 2014); mouse downregulated in Rfx3 −/− mutant embryos (Lemeille et al., 2020); human motile cilia signature (Patir et al., 2020); (3) Localization: trypanosome basal bodies or flagellum localization (Dean et al., 2017), from http://tryptag.org.

**Table S3: Full details of primary and secondary screens in *Drosophila* and *C. elegans*.** (**A**) Datasheet summarizing results of primary screen for each ciliary cluster gene conserved in *C. elegans* and *Drosophila* (non-conserved genes included for ease of comparison with other tables). For *C. elegans*, putative loss of function mutants were obtained and screened using the dye-fill assay. Phenotypes in amphid (head) and phasmid (tail) neurons were scored separately on a scale from 0 (no phenotype) to 4 (dye-fill null) and the stronger of the two reported as the overall score in Fig. 3 (see table for definitions). For *Drosophila*, multiple RNAi lines (where available) were obtained and scored for uncoordination, male fertility and viability (as applicable) with each of the three drivers, Elav for depletion in sensory neurons, Nanos and Bam combined for germline depletion, and Tubulin for ubiquitous depletion. Phenotypes were scored on a scale from 0 (no phenotype) to 4 (maximum phenotype) and the strongest neuronal and male fertility phenotype reported in Fig. 3 (see table for definitions). Controls included in the primary screen are shown on a separate datasheet. Also included on that datasheet are the phenotypic data obtained in *Drosophila* for additional CFAP97 and CFAP299-related genes. (**B**) Datasheets summarizing results of secondary screen in *Drosophila*, with separate sheets for chordotonal neurons and sperm. In each case a single RNAi line from the primary screen was chosen for further phenotypic characterization. For chordotonal neurons, phenotypes were scored by DIC (number of cilia/scolopidium, number of morphologically defective cilia, and for selected genes position of ciliary dilation relative to the cap cell) and NompC/Sas-4 staining (percentage of defective cilia per scolopidium). Plotted in Fig. 4C are the number of intact cilia based on DIC and NompC staining (generally a lower number, given that this assay is more sensitive). For sperm, the severity of defects at the six distinct stages of spermatogenesis is recorded on a scale from 0 (no phenotype) to 4 (maximum phenotype), as well as the motility of sperm stored in the seminal vesicle. Plotted in Fig. 4G is the stage at which phenotypes of severity 2 or more are first observed, as well as the stage at which the maximum phenotype of 4 is reached.

## Notes

### Competing Interest Statement

The authors have declared no competing interest.

